# A detergent-free workflow for native membrane proteomics using Peptergents

**DOI:** 10.64898/2026.08.12.744532

**Authors:** Frank Antony, Ashim Bhattacharya, Hiroyuki Aoki, Mohan Babu, Franck Duong van Hoa

**Affiliations:** Department of Biochemistry and Molecular Biology, Faculty of Medicine, Life Sciences Institute, University of British Columbia, Vancouver, British Columbia, Canada; Department of Biochemistry, Faculty of Science, University of Regina, Regina, Saskatchewan, Canada

**Keywords:** Peptergent, Detergent-free extraction, Thermal proteome profiling, Membrane Proteomics, Ligand binding, Membrane mimetics, Native membrane

## Abstract

Quantitative membrane proteomics remains fundamentally limited by sample preparation because detergent extraction can perturb membrane protein interactions, ligand-responsive conformations, and higher-order assemblies before mass spectrometric analysis. Here, we demonstrate that peptide-based surfactants (Peptergents) enable a complete detergent-free workflow for native membrane proteomics. Membrane proteins are extracted directly from biological membranes while preserving their structural and functional integrity and remaining fully compatible with downstream LC–MS/MS workflows. Functional preservation is evidenced by maintenance of ligand-responsive conformations in the ABC transporter MsbA and the endogenous GPCR P2RY12, together with stabilization of the detergent-sensitive nine-subunit holo-translocon HTL, indicating that fragile membrane protein assemblies remain intact. At the proteome level, despite recovering fewer membrane proteins than conventional detergent extraction, Peptergent consistently generates higher peptide signal intensities, retains tissue-specific membrane proteome signatures, and preferentially enriches endoplasmic reticulum-associated metabolic networks, including cytochrome P450 enzymes and their interaction network. Together, these findings establish Peptergents as a broadly applicable membrane extraction technology for LC–MS/MS-based membrane proteomics, preserving native membrane organization and expanding the proteomics toolbox for biochemical, structural, and systems-level analyses of membrane proteins.

**In Brief Statement:** This study establishes Peptergents as a detergent-free membrane extraction technology for LC–MS/MS-based membrane proteomics. Peptergent extraction preserves ligand-responsive membrane proteins, fragile membrane protein assemblies, and tissue-specific membrane proteome signatures while remaining fully compatible with quantitative proteomic workflows. These findings provide a broadly applicable strategy for preserving native membrane organization for biochemical, structural, and systems-level analyses of membrane proteins.

**Graphical Abstract:** 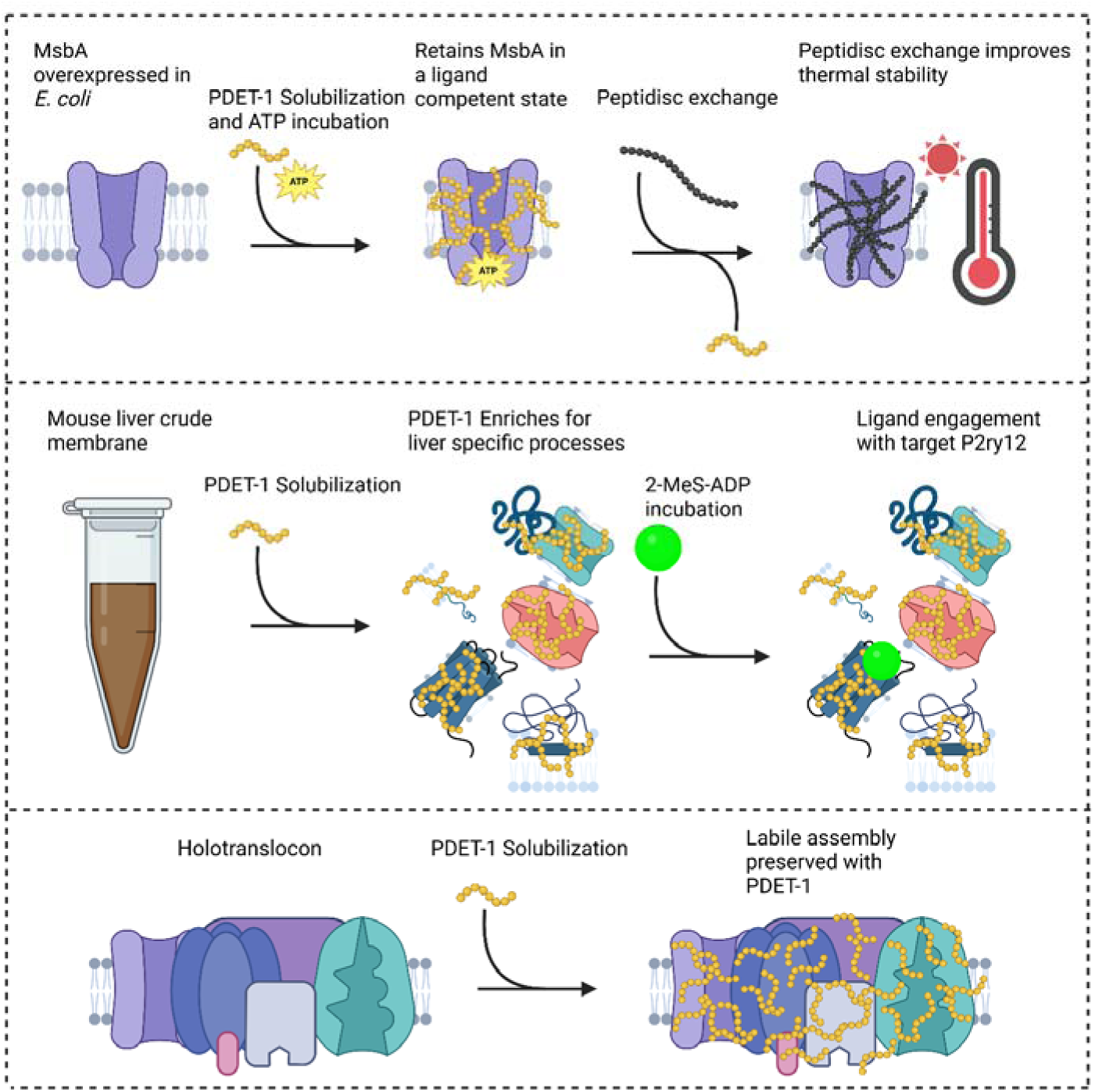

**Highlights:**

- Peptergents preserve ligand-responsive membrane proteins.
- Support chemoproteomics in thermal proteome profiling assays.
- Simplify membrane proteomics workflow.
- Maintain native tissue-specific membrane biology.
- Preserve fragile membrane protein assemblies.

## Introduction

Integral membrane proteins (IMPs) constitute approximately 30% of the human proteome^1^ and mediate virtually every aspect of cellular communication, including signaling, transport, metabolism, and cell–cell interactions^2^. They also represent the largest class of therapeutic drug targets^3^. Despite major advances in mass spectrometry, quantitative proteomics, and computational analysis, membrane proteins remain among the least accessible components of the proteome because the biological information captured in membrane proteomic datasets remains strongly influenced by sample preparation^4^.

Conventional detergents efficiently solubilize biological membranes by disrupting the lipid bilayer and replacing the native membrane environment with detergent micelles. Although highly effective at membrane solubilization while maintaining many membrane proteins in functional states, detergent extraction can destabilize membrane proteins, dissociate higher-order assemblies, disrupt protein–lipid interactions, alter ligand-responsive conformations, and ultimately reduce the biological fidelity of membrane proteomic datasets before LC–MS/MS analysis even begins^4–6^. Furthermore, because most detergents are incompatible with mass spectrometry, they must be removed prior to analysis, introducing additional processing steps that can reduce protein recovery and increase experimental variability^7^.

These limitations have motivated the development of detergent-free membrane mimetics, including amphipols^8^, styrene-maleic acid lipid particles^9^, nanodiscs^10^, and peptide-based membrane mimetics such as Salipro^11^ and Peptidiscs^12–15^. These technologies have transformed structural biology and membrane protein biochemistry by stabilizing membrane proteins in native-like environments after extraction. However, most still rely on an initial detergent solubilization step or require removal of the membrane mimetic before LC–MS/MS analysis^7^. As a result, membrane extraction and downstream proteomic analysis remain largely independent stages of the workflow, and native membrane organization may already be altered before quantitative analysis begins.

Peptide-based membrane extraction reagents (Peptergents) directly solubilize biological membranes without detergents^16,17^. Previous studies have shown that amphiphilic peptides can extract and stabilize selected membrane proteins for biochemical and structural applications, establishing peptide-mediated extraction as a viable alternative to detergent solubilization for individual membrane proteins^18–20^. Whether this strategy can be extended to preserve the native organization of complex membrane proteomes, however, has remained unknown. In particular, it is unclear whether detergent-free extraction can simultaneously preserve ligand-responsive membrane proteins, fragile endogenous membrane assemblies, tissue-specific membrane biology, and compatibility with LC–MS/MS-based membrane proteomic workflows.

Here, we demonstrate that the Peptergent PDET-1 enables a detergent-free workflow for native membrane proteomics. Using complementary biochemical, functional, and quantitative proteomic analyses spanning purified membrane proteins, native membrane proteomes, and endogenous membrane protein complexes, we show that detergent-free extraction preserves ligand-responsive conformations, stabilizes fragile membrane assemblies, maintains tissue-specific membrane biology, and integrates seamlessly with LC–MS/MS workflows. Together, these findings establish Peptergents as a broadly applicable detergent-free membrane extraction strategy that preserves native membrane organization while enabling LC–MS/MS-based membrane proteomics.

## Results

### PDET-1 extracts MsbA into an exchangeable peptide–protein assembly

We first evaluated whether PDET-1 could directly extract the homodimeric ABC transporter MsbA from *E. coli* membranes. Crude membranes containing overexpressed His-tagged MsbA were incubated with increasing concentrations of PDET-1, followed by Ni-NTA affinity purification. MsbA recovery increased with PDET-1 concentration, reaching a maximum at 2 mg/mL before plateauing. At this concentration, PDET-1 recovered approximately 40% of the MsbA obtained by extraction with 1% DDM while achieving comparable purity (**Figure 1A-C**).

**Figure 1.**
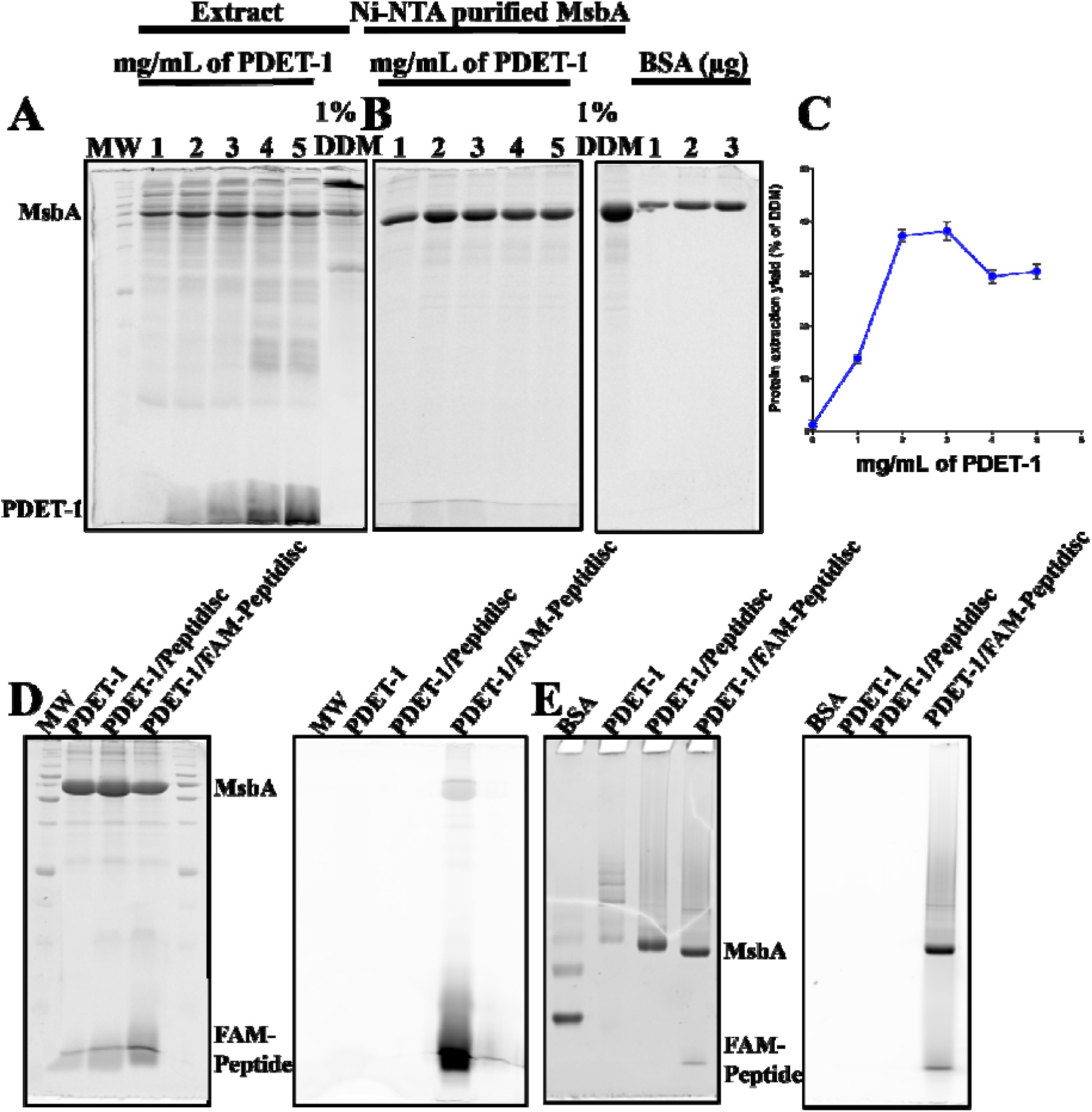
Extraction of MsbA from *E. coli* membranes with PDET-1 and subsequent exchange into Peptidisc. **(A)** SDS-PAGE analysis of extracts from His-tagged MsbA-enriched membranes following incubation with the indicated concentrations of PDET-1. The 1% DDM extract is shown in the rightmost lane as a reference, and molecular weight markers are shown in the leftmost lane. PDET-1 peptides migrate near the bottom of the gel as low-molecular-weight bands. Samples were not heat-denatured before SDS-PAGE; consequently, a fraction of MsbA in the crude DDM extract migrates as a high-molecular-weight species. **(B)** His-tagged MsbA was purified from the extracts shown in (A) by Ni-NTA affinity chromatography, and the resulting eluates were analyzed by SDS-PAGE. Bovine serum albumin (BSA) is loaded on the same gel as a protein quantity standard. **(C)** Relative recovery of purified MsbA following extraction with the indicated concentrations of PDET-1. The amount recovered with 1% DDM was defined as 100%. Values represent the mean ± SD of three independent experiments. Quantified using ImageJ densitometry of the MsbA band. **(D)** SDS-PAGE analysis of purified MsbA in PDET-1 alone, or after exchange into unlabeled Peptidisc (PDET-1/Peptidisc), or FAM-labeled Peptidisc (PDET-1/FAM-Peptidisc). The left and right panels show Coomassie staining and fluorescence scan of the same gel. Under denaturing conditions, the FAM-labeled Peptidisc dissociates from MsbA and migrates as free peptide. **(E)** Native-PAGE analysis of the samples prepared in (D). BSA (66 kDa) is loaded on the same gel as a reference marker. The left and right panels show Coomassie staining and fluorescence scan of the same gel.

We next examined the oligomeric state of PDET-1-extracted MsbA. The SDS-PAGE analysis showed a single purified MsbA species, whereas native PAGE revealed a ladder of discrete bands, indicating that the transporter was solubilized as a heterogeneous population of peptide–protein assemblies. Exchanging PDET-1 with the Peptidisc scaffold during purification converted this heterogeneous population into a single, well-defined species on native PAGE, consistent with formation of a monodisperse Peptidisc-stabilized MsbA dimer. Efficient replacement of PDET-1 by the Peptidisc scaffold was confirmed using a fluorescently labeled (FAM) Peptidisc peptide (**Figure 1D-E**).

Together, these results demonstrate that PDET-1 extracts IMPs into a soluble peptide–protein assembly that combines membrane extraction with protein stabilization while remaining readily exchangeable for downstream applications.

### PDET-1 preserves the ligand-responsive conformation of MsbA

Having shown that PDET-1 can solubilize MsbA, we next asked whether this extraction process also preserves the protein in a functional, ligand-competent state. During ATP hydrolysis, orthovanadate (Vi) forms a stable ADP–Vi complex that mimics the transition state of phosphate release, trapping MsbA in its ATP-hydrolysis transition-state conformation and thereby increasing its thermal stability^6^. Ligand-dependent thermal stabilization therefore provides a functional readout of ATP binding, hydrolysis competence, and protein conformational integrity.

MsbA prepared by PDET-1 extraction, PDET-1 followed by Peptidisc exchange, or DDM solubilization was subjected to ATP-Vi-dependent thermal denaturation. In the absence of ATP-Vi, MsbA denatures at 45°C following DDM solubilization, 48°C after PDET-1 extraction, and 51°C following PDET-1/Peptidisc exchange. Addition of ATP-Vi produced a clear rightward shift in the denaturation temperature of PDET-1-extracted MsbA (48°C to 51°C) and an even larger shift following Peptidisc exchange (51°C to 57°C). In contrast, DDM-solubilized MsbA exhibited no detectable ATP-Vi-dependent stabilization (**Figure 2**; **Full gel lanes Supplemental Figures 1-2**).

**Figure 2.**
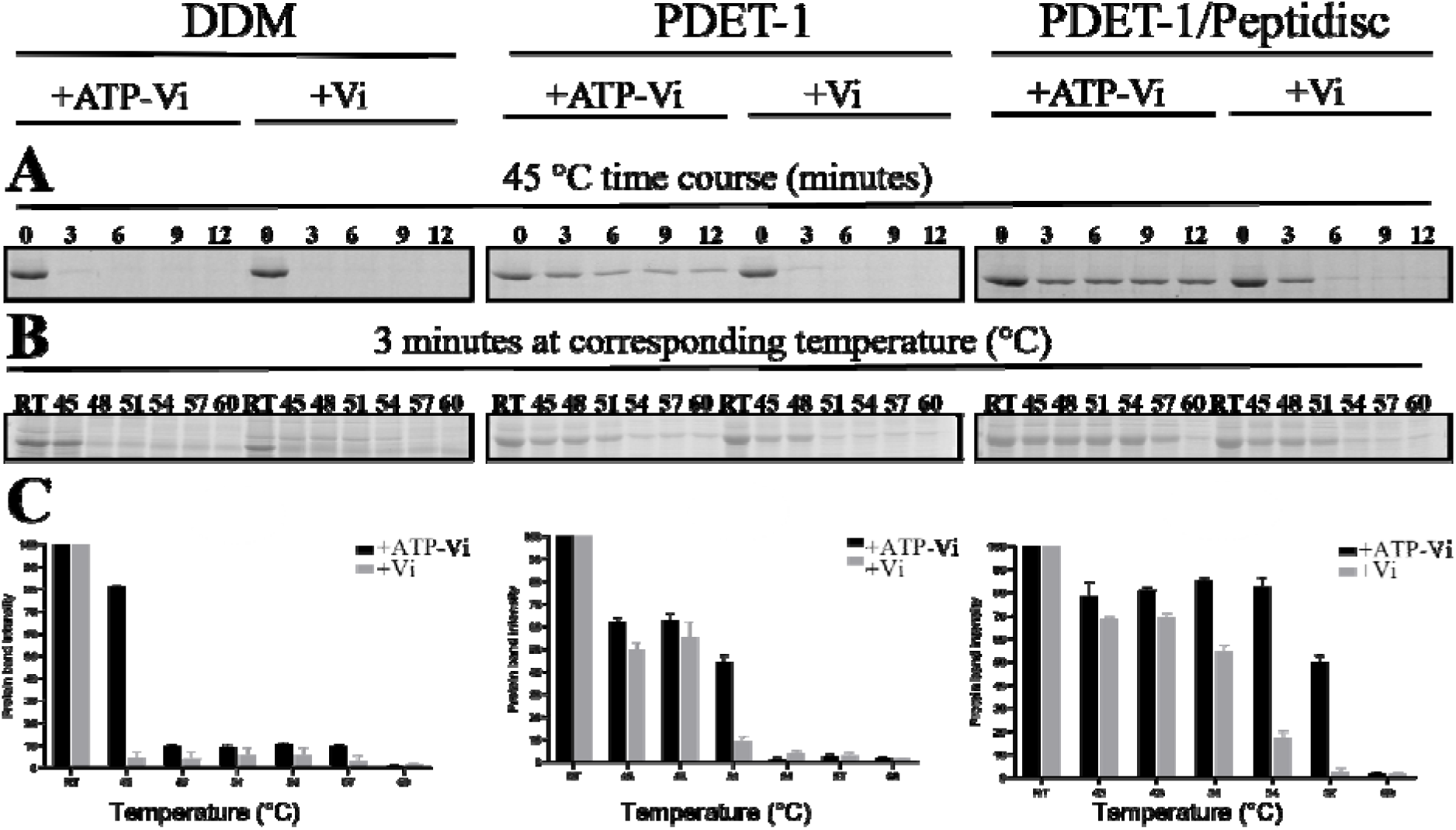
Ligand-dependent thermal stabilization of MsbA following solubilization with DDM, PDET-1, or PDET-1/Peptidisc. **(A)** SDS-PAGE analysis of soluble MsbA remaining after ATP-Vi-dependent thermal denaturation following extraction with DDM, PDET-1, or PDET-1 followed by exchange into Peptidisc. Samples were incubated in the absence or presence of ATP-Vi at 45 °C for the indicated time, before centrifugation and analysis of the soluble fraction by SDS-PAGE. The full lane gel can be found in supplemental figure 1. **(B)** Representative SDS-PAGE analysis of soluble MsbA crude membrane remaining after ATP-Vi-dependent thermal denaturation following extraction with DDM, PDET-1, or PDET-1 followed by exchange into Peptidisc. Samples were incubated in the absence or presence of ATP-Vi at the indicated temperature for 3 minutes, before centrifugation and analysis of the soluble fraction by SDS-PAGE. Experiment was performed in triplicate and the full lane gels can be found in supplemental figure 2. (**C**) Quantification of soluble MsbA remaining after thermal denaturation as done in (B). Band intensities were quantified by densitometry using ImageJ on the MsbA band, normalized to the unheated sample (RT = 100%), and plotted as a function of temperature. Black symbols, +ATP-Vi; gray symbols, −ATP-Vi. Values represent the mean ± SD of three independent experiments.

These observations establish that PDET-1 preserves the ligand-responsive conformation of MsbA following membrane extraction. Subsequent exchange into Peptidisc further enhances thermal stability while maintaining ligand responsiveness, whereas conventional DDM solubilization abolishes detectable ligand-dependent stabilization.

### Peptergent enables ligand-dependent thermal proteome profiling of the native membrane proteome

Having established that Peptergent preserves ligand binding for the purified transporter MsbA, we next asked whether this behavior extends to the native membrane proteome using membrane-mimetic thermal proteome profiling (MM-TPP). MM-TPP detects ligand-induced thermal stabilization across the entire proteome by LC-MS/MS and therefore provides a stringent test of whether receptor–ligand interactions are preserved following membrane extraction^6^. We used the selective P2RY12 agonist 2-methylthio-ADP (2-MeS-ADP)^21^, reasoning that successful preservation of receptor function should identify P2RY12 as the most significantly stabilized protein in the membrane proteome.

Consistent with this expectation, P2RY12 was the most significantly stabilized protein in both Peptergent-based workflows, including PDET-1 alone (−log□□*p* = 5.01, log□FC = 2.20) and PDET-1 followed by exchange into Peptidisc (−log□□*p* = 6.51, log□FC = 4.00). The stronger stabilization of P2RY12 obtained with Peptidisc mirrors the enhanced thermal stability previously observed for MsbA, suggesting that Peptidisc better stabilizes proteins after their initial Peptergent extraction. In contrast, DDM-solubilized samples showed no significant stabilization of P2RY12 (−log *p* = 1.28, log FC = −0.25) (**Figure 3**).

**Figure 3.**
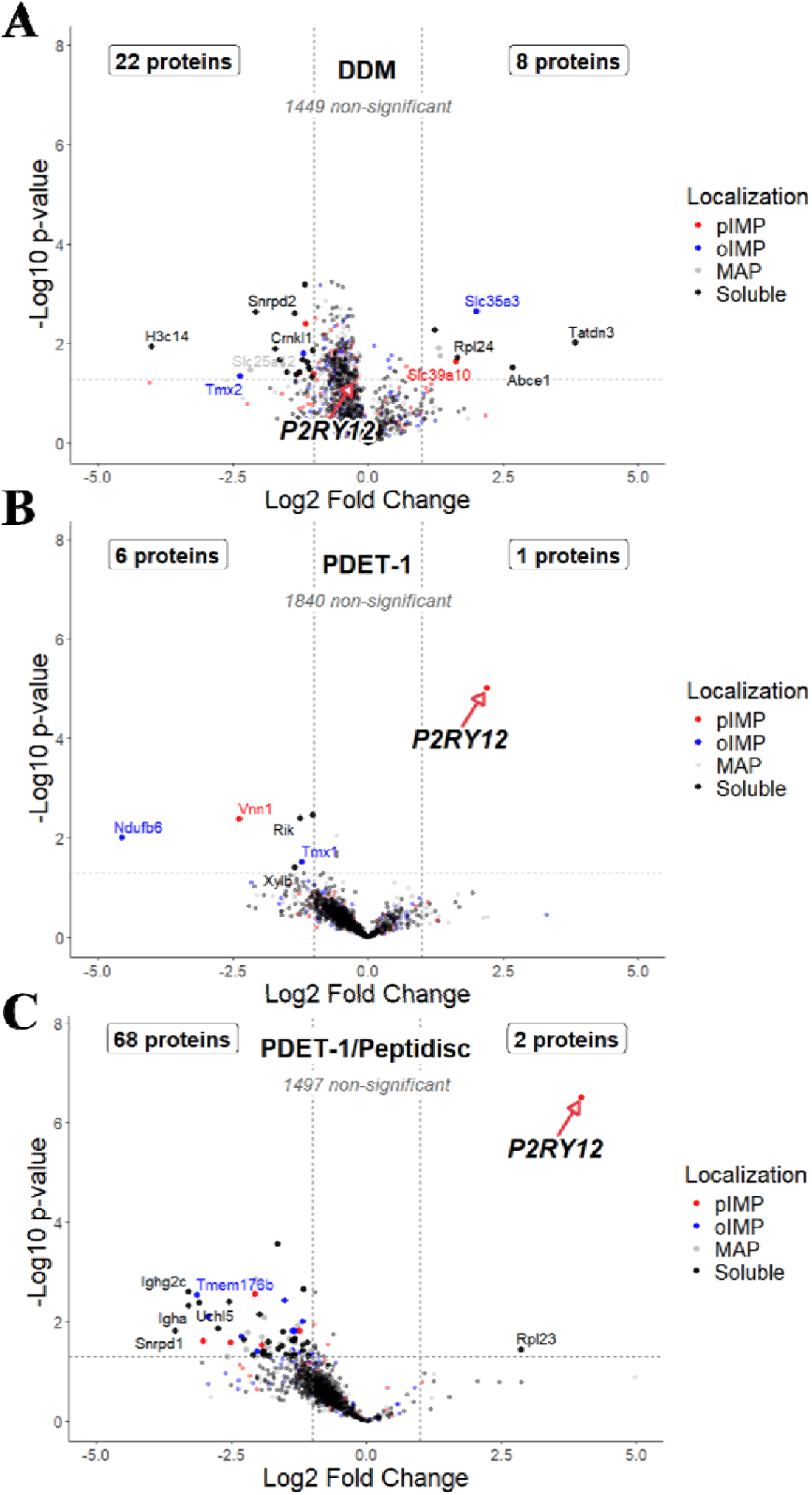
Membrane mimetic-thermal proteome profiling analysis of mouse liver membranes solubilized with DDM, PDET-1 or PDET-1/Peptidisc and incubated with 2-MeS-ADP. **(A–C)** Volcano plots showing differential thermal stability of mouse liver membrane proteins following treatment with 2-MeS-ADP relative to vehicle control after solubilization with (A) DDM, (B) PDET-1, or (C) PDET-1 exchanged into Peptidisc. Log□ fold change (2-MeS-ADP versus vehicle) is plotted against −log□□(p). Dashed vertical and horizontal lines indicate the fold-change and significance thresholds (p < 0.05), respectively. Proteins are colored according to localization: plasma membrane integral membrane proteins (pIMP, red), organellar integral membrane proteins (oIMP, blue), membrane-associated proteins (MAP, gray), and soluble proteins (black). P2RY12 is highlighted in each panel (red arrow); log□fold change and −log (p) values were, respectively: DDM, −0.25 and 1.28 (not significant); PDET-1, 2.20 and 5.01; PDET-1/Peptidisc, 3.99 and 6.51. Insets summarize the numbers of significantly stabilized and destabilized proteins in each localization category.

Collectively, these MM-TPP results demonstrate that Peptergent preserves native receptor–ligand interactions not only for purified membrane proteins but also within complex native membrane proteomes, thereby enabling functional membrane proteomics. While direct PDET-1 extraction is sufficient to detect ligand-dependent stabilization, subsequent exchange into Peptidisc further enhances the thermal response, whereas conventional DDM solubilization abolishes detectable ligand-dependent stabilization.

### PDET-1 preserves membrane proteome composition despite reduced protein coverage

Having established that PDET-1 preserves membrane proteins in a ligand-competent state, we next evaluated its suitability for membrane proteome profiling using mouse liver membranes, chosen for their complex and differentiated membrane composition. PDET-1 extraction conditions were first optimized by SDS-PAGE, with recovery increasing up to 2 mg/mL before reaching a plateau. At this concentration, SDS-PAGE analysis indicated that PDET-1 achieved recovery comparable to DDM (**Supplemental Figure 3**). Next, we compared the proteomes recovered by PDET-1 and DDM using bottom-up LC-MS/MS with three biological replicates. Proteins were classified as soluble proteins (Sol), membrane-associated proteins (MAPs), plasma membrane integral membrane proteins (pIMPs), or organellar integral membrane proteins (oIMPs), with pIMPs and oIMPs collectively designated integral membrane proteins (IMPs). Consistent with the strong membrane-disrupting properties of conventional detergents, DDM identified more proteins overall than PDET-1 (2633 ± 8 versus 1944 ± 22), including more integral membrane proteins (606 ± 2 versus 479 ± 1), reflecting greater proteome coverage. Despite this difference in overall recovery, the proportion of tIMPs was nearly identical between the two extraction methods (23–25%; **Table 1**).

**Table 1.** Proteome coverage and subcellular distribution of mouse liver membrane proteins solubilized with DDM or PDET-1. Total proteins identified by MaxQuant LFQ intensity DDA-MS following solubilization of mouse liver membranes with DDM or PDET-1. Proteins were classified as soluble, membrane-associated (MAP), total integral membrane proteins (tIMP), organellar integral membrane proteins (oIMP), or plasma membrane integral membrane proteins (pIMP) based on Phobius topology annotation. Values represent the mean ± SD across biological triplicate; percentages in parentheses indicate the proportion of each category relative to the total number of proteins identified for each extraction condition.

| MS sample<br>(mean ± SD) | TPs | | | | IMPs | |
| --- | --- | --- | --- | --- | --- | --- |
|  | Total proteins<br>(TPs) | Soluble proteins<br>(SPs) | Membrane associated proteins<br>(MAPs) | Integral membrane proteins<br>(IMPs) | pIMPs | oIMPs |
| mLiver DDM | 2633 ± 8 | 1234 ± 4<br>(46.9%) | 1399 ± 10<br>(53.1%) | 606 ± 2<br>(23.0%) | 262 ± 3<br>(10.0%) | 344 ± 3<br>(13.1%) |
| mLiver PDET-1 | 1944 ± 22 | 898 ± 18<br>(46.2%) | 1046 ± 18<br>(53.8%) | 479 ± 1<br>(24.6%) | 193 ± 3<br>(9.9%) | 286 ± 3<br>(14.7%) |

To determine whether the lower protein recovery by PDET-1 affected the composition of the recovered membrane proteome, we compared the subcellular localization of the 200 most abundant proteins identified by each extraction method. Despite recovering fewer proteins overall, PDET-1 produced a membrane proteome composition that closely matched DDM, with no localization category differing by more than three proteins (**Supplemental Figure 4**).

Similarly, the distributions of transmembrane segment numbers among IMPs and the functional classes represented within pIMPs were indistinguishable between the two methods (**Supplemental Figure 5A-B**).

Together, these analyses show that although DDM identifies a larger number of membrane proteins than PDET-1, both workflows recover membrane proteomes with highly similar structural and functional composition. Thus, the principal difference lies in proteome depth rather than in selective representation of particular membrane protein classes.

### DDM and PDET-1 exhibit distinct protein recovery profiles

The preceding analyses addressed qualitative proteome composition. We here examined quantitative recovery, asking whether individual membrane proteins differed in abundance between the two extraction workflows. Of the 2,469 proteins detected across both datasets, 419 were significantly enriched by PDET-1 and 437 by DDM, whereas the remaining 1,613 proteins (65.3%) showed no significant difference (**Figure 5A**).

To determine whether these enrichment biases were localized to specific membrane compartments, we examined protein localization across seven membrane-associated compartments; only endoplasmic reticulum (ER) proteins showed a significant overall shift toward PDET-1 enrichment (n = 480, median log2FC = 0.53, adjusted p = 4.64 × 10) (**Figure 4B**). Within this ER subset, 132 proteins were more abundant following PDET-1 extraction compared with 58 for DDM, and most proteins favored by PDET-1 were IMPs (79 of 132; 60%) (**Supplemental Figure 5C**). These proteins were strongly associated with hepatic metabolic functions. Of the 37 cytochrome P450 (CYP) family members detected, 26 (70%) showed significantly higher abundance with PDET-1, whereas none favored DDM. Beyond individual CYP450 enzymes, PDET-1 also preferentially recovered their obligate redox partner NADPH-cytochrome P450 reductase^22^ (Por; log2FC = 1.25, *p* < 0.001) (**Supplemental Figure 6A**). Likewise, high-confidence CYP-interacting proteins (STRING combined score ≥700)^23^ showed significantly greater enrichment by PDET-1 than the remainder of the quantified proteome (**Supplemental Figure 6B**). Consistent with this observation, TissueEnrich analysis, which classifies proteins as tissue-enriched based on GTEx human transcriptomic data^24^, showed that liver-enriched marker proteins were significantly overrepresented among PDET-1-enriched proteins compared with DDM-enriched proteins (**Figure 4C**). Together, these findings indicate that PDET-1 preferentially recovers the CYP450 metabolic network rather than isolated enzymes.

**Figure 4.**
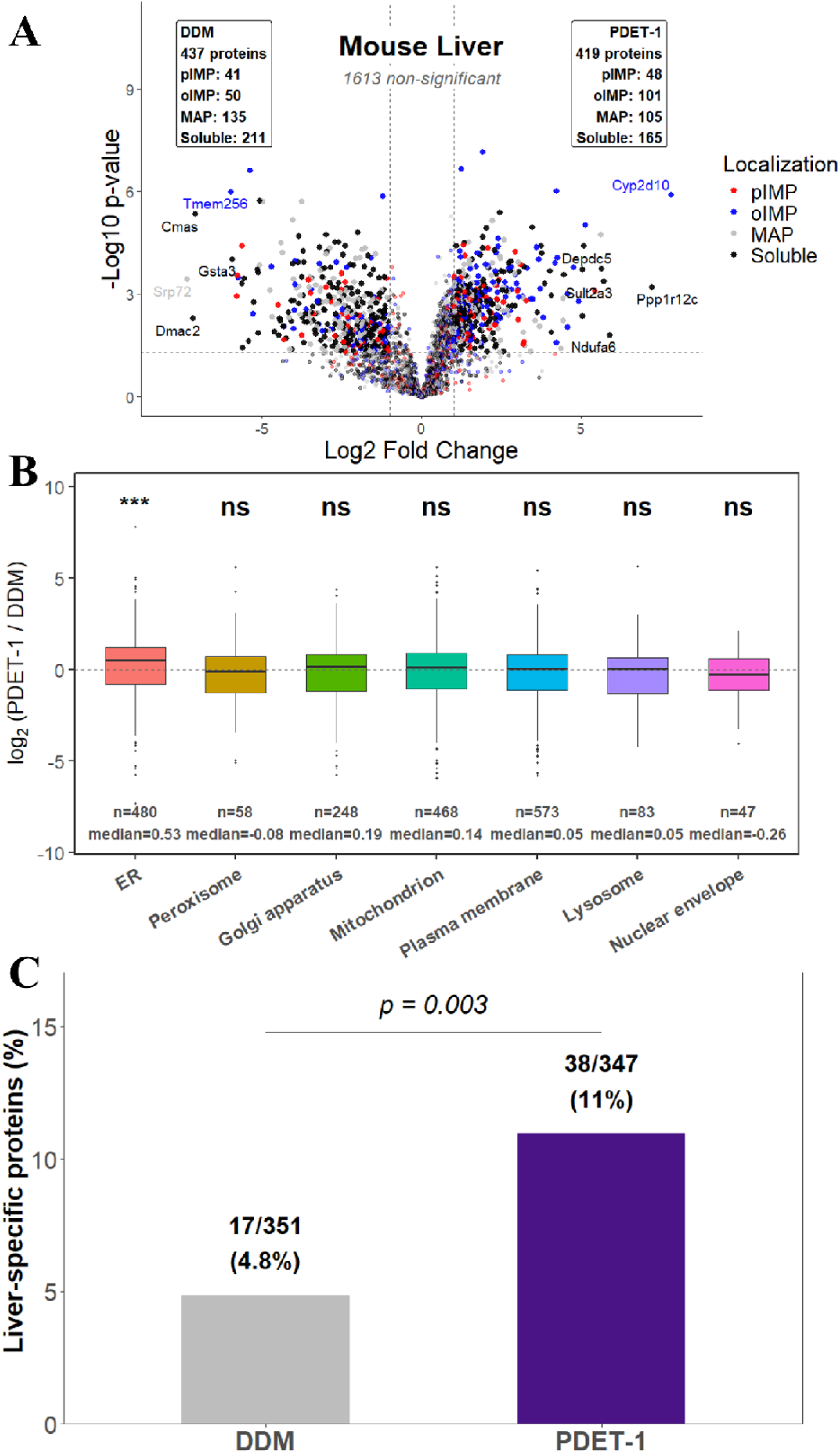
Comparative analysis of mouse liver membrane protein recovery following extraction with DDM or PDET-1. (**A**) Volcano plot comparing protein abundance following extraction with DDM or PDET-1. Log□fold change is plotted against −log□□(p) calculated as a two-tailed t-test. (**B**) Distribution of log□ fold change values for proteins assigned to the indicated membrane compartments based on Gene Ontology cellular component annotations. Boxes indicate the median and interquartile range; whiskers denote 1.5 × the interquartile range. Statistical significance was assessed using a one-sample Wilcoxon signed-rank test with Benjamini–Hochberg correction (ER: p = 6.6 × 10□□, p.adj = 4.6 × 10□□; Peroxisome: p = 0.491, p.adj = 0.573; Golgi apparatus: p = 0.429, p.adj = 0.573; Mitochondrion: p = 0.975, p.adj = 0.975; Plasma membrane: p = 0.142, p.adj = 0.336; Lysosome: p = 0.310, p.adj = 0.543; Nuclear envelope: p = 0.144, p.adj = 0.336). **, adjusted p < 0.001; ns, not significant. **(C)** Percentage of significantly enriched proteins annotated as liver-specific markers in the DDM- and PDET-1-enriched datasets. Liver-specific proteins were defined using the TissueEnrich GTEx-derived tissue-enriched gene set. Numbers above the bars indicate the number of liver-specific proteins relative to the total number of significantly enriched proteins (DDM: 17/351, 4.8%; PDET-1: 38/347, 11%). Statistical significance was assessed using Fisher’s exact test (odds ratio = 2.41, 95% CI: 1.30–4.66, p = 0.003).

By comparison, among the 437 DDM-enriched proteins, 41 were annotated with the Gene Ontology term *translation*. This enrichment was driven primarily by components of the 40S ribosomal subunit and translation initiation complexes. These observations indicate distinct recovery biases, with PDET-1 preferentially enriching ER-associated metabolic networks and DDM preferentially recovering translation initiation machinery rather than ribosomal proteins more broadly.

### PDET-1 yields higher quantitative peptide signals despite reduced peptide coverage

We next compared peptide recovery and quantitative performance between the DDM and PDET-1 workflows. DDM identified substantially more peptides than PDET-1 (21,750 versus 12,660 detected in at least two of three biological replicates), whereas peptides detected by PDET-1 were modestly more hydrophobic (**Figure 5A**). Despite recovering fewer peptides, PDET-1 yielded significantly higher signal intensities overall, resulting in a 34.9% greater summed peptide intensity (**Figure 5B**).

**Figure 5.**
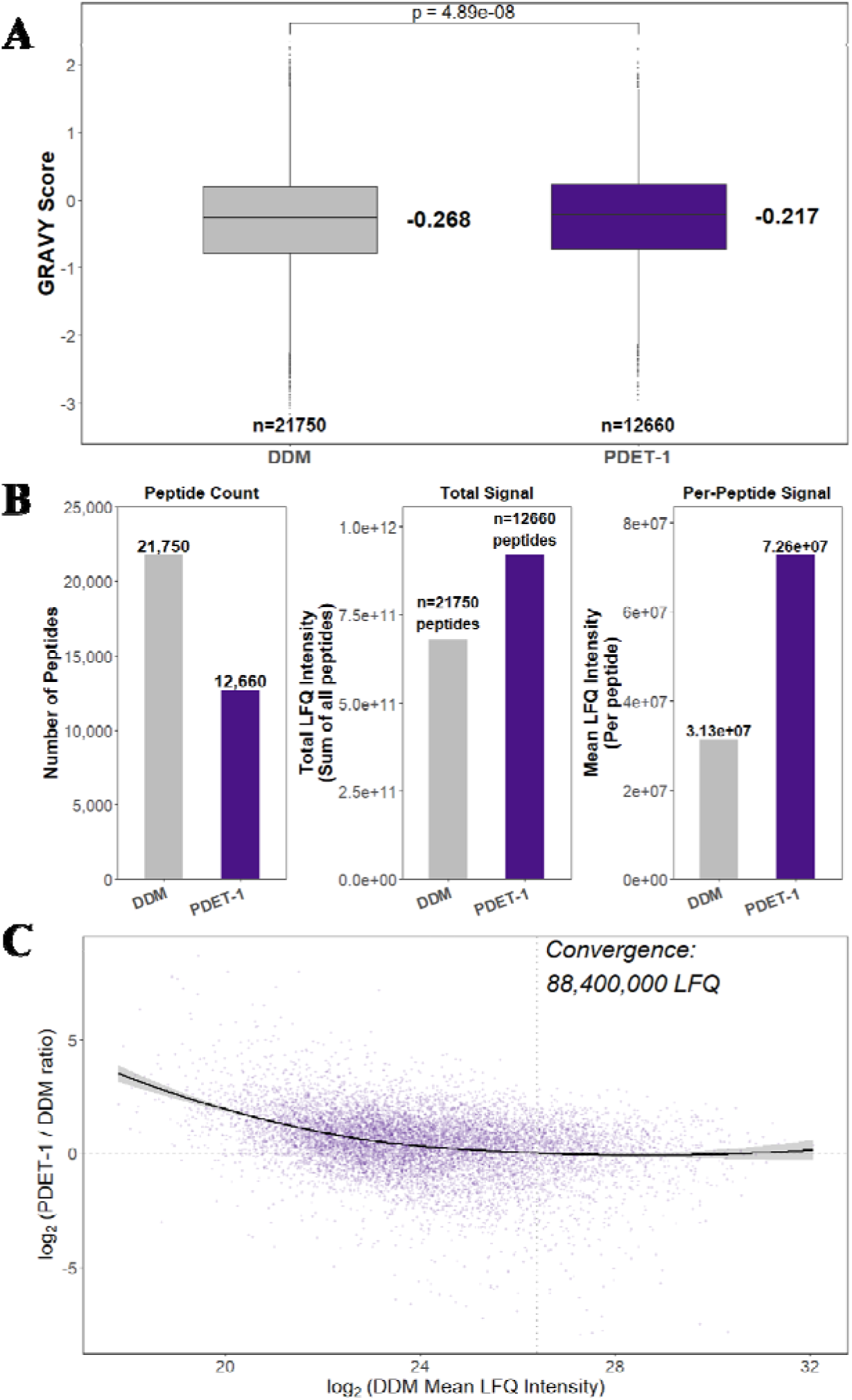
Comparative peptide properties and recovery following extraction with DDM or PDET-1. **(A)** Distribution of peptide GRAVY (grand average of hydropathicity) scores identified following extraction with DDM or PDET-1 (DDM: n = 21,750, median = −0.268; PDET-1: n = 12,660, median = −0.217). Statistical significance was assessed using a two-sided Wilcoxon rank-sum test with continuity correction (W = 132,829,839, p = 4.89 × 10□□). (**B**) Comparison of peptide count (left), total LFQ intensity (middle), and mean LFQ intensity per peptide (right) between DDM and PDET-1. **(C)** Scatter plot of log (PDET-1/DDM) peptide abundance versus mean log LFQ intensity for peptides identified in both extraction conditions (n = 10,703 shared peptides). Log-transformed DDM and PDET-1 intensities were strongly correlated (Pearson’s r = 0.784, 95% CI: 0.777–0.791, p < 2.2 × 10□¹□). The black line represents a LOESS fit (span = 0.75, degree = 2; residual standard error = 1.29) with the shaded region indicating the 95% confidence interval. The fit crosses zero (equal DDM/PDET-1 abundance) at a mean DDM intensity of ∼8.8 × 10□ LFQ units, indicating convergence of the two methods’ sensitivity above this intensity threshold.

To determine the basis of this quantitative difference, we focused on the 10,703 peptides detected by both workflows. Despite identifying more peptides overall, DDM yielded lower signal intensities for peptides shared between the two datasets than PDET-1. This advantage was greatest among low-abundance peptides and progressively diminished at higher intensities until both methods converged (**Figure 5C**). Conversely, 99.1% of peptides detected exclusively by DDM fell below the empirical convergence point, indicating that DDM’s additional peptide coverage consisted almost entirely of very low-intensity peptides (**Supplemental Figure 7A**).

The PDET-1 intensity advantage was independent of transmembrane segment number and amino acid composition (**Supplemental Figure 7B-C**), but progressively declined with increasing peptide hydrophobicity and was absent for the most hydrophobic transmembrane peptides (**Supplemental Figure 8**). These findings indicate that peptide physicochemical properties contribute, at least in part, to the quantitative differences observed between the two workflows.

Consistent with the protein-level analysis, PDET-1 CYP-derived peptides (425 peptides representing 26 genes) exhibited significantly greater signal intensity than the remaining shared peptide population, accounting for 7.7% of the overall intensity difference despite representing only 4% of shared peptides. In contrast, peptides detected exclusively by DDM showed no comparable biological enrichment, indicating that DDM’s additional peptide coverage primarily reflects non-specific recovery of low-abundance peptides rather than preferential enrichment of a specific biological pathway.

Together, these findings are consistent with the observed differences between the two workflows. DDM extends peptide coverage primarily through detection of very low-abundance peptides, whereas PDET-1 yields substantially stronger quantitative signals from the peptides it recovers. Because the two workflows differ in both membrane extraction and downstream sample preparation, the relative contribution of each step cannot be distinguished from the present data.

### Differences in extraction workflow do not compromise recovery of tissue-specific biology

We next asked whether the extraction workflow influences recovery of tissue-specific biological information by comparing mouse liver and brain membrane proteomes. As observed for liver, DDM identified more proteins than PDET-1 in brain (2984 ± 19 versus 2071 ± 72), whereas PDET-1 again recovered a modestly higher proportion of IMPs (26.2% versus 24.8%) (**Supplemental Table 1**). Principal component analysis of the mouse liver and brain datasets identified tissue identity as the dominant source of variation (PC1, 68.3% of total variance), substantially exceeding the contribution of the extraction method (PC2, 16.9%) (**Supplemental Figure 9A**). Consistent with this, both workflows clearly separated liver and brain proteomes and recovered extensive sets of tissue-enriched and tissue-exclusive proteins (**Supplemental Figure 9B-C**).

To determine whether these differences reflected authentic tissue biology rather than workflow-specific bias, we validated the tissue-enriched protein sets using three independent approaches. TissueEnrich confirmed significant enrichment of canonical liver and cerebral cortex markers for both methods, although a higher proportion of PDET-1-enriched proteins were cerebral cortex markers (20.4% versus 11.6%; **Supplemental Figure 10**). Gene Ontology analysis identified the expected biological programs for each tissue, including xenobiotic and lipid metabolism in liver and neuronal processes in the brain (**Supplemental Figure 11A-B**).

Likewise, comparison with published organ-specific plasma membrane protein datasets^25^ showed that both workflows recovered the majority of established liver- and brain-specific markers (**Supplemental Figure 11C**).

Together, these findings show that although DDM and PDET-1 differ in extraction efficiency and quantitative recovery of specific protein classes, both faithfully preserve the biological signatures that distinguish tissues.

### PDET-1 preserves labile membrane protein complexes

Having established that PDET-1 preserves membrane proteins in a ligand-competent state and recovers biologically coherent membrane proteomes, we next asked whether it also preserves labile membrane protein complexes. The coordinated recovery of CYP450 enzymes together with Por and their associated interaction network suggested that PDET-1 may preserve native membrane protein assemblies rather than extracting proteins as isolated entities. To test this directly, we examined the bacterial holo-translocon (HTL), a ∼250-kDa membrane protein insertion and translocation machine composed of nine subunits spanning 31 transmembrane helices^26^. The HTL comprises the SecYEG protein-conducting channel, the SecDF-YajC complex, the YidC membrane insertase, and the single-pass membrane chaperones PpiD and YfgM^27^. Because interactions between these modules are relatively weak, the intact complex readily dissociates during DDM solubilization into smaller subcomplexes, preventing recovery of the complete assembly by affinity purification.

Using PDET-1, affinity purification of His-tagged HTL recovered all nine HTL subunits above the BL21 background, including the SecYEG core complex, the SecDF-YajC module, YidC, and the non-overexpressed ancillary proteins PpiD and YfgM (**Figure 6A**). Furthermore, these findings were reproduced under native expression conditions. Affinity purification of chromosomally SPA-tagged SecY recovered the complete HTL together with YfgM and PpiD relative to the DY330 control (**Figure 6B**), demonstrating that preservation of the intact holo-translocon is not an artifact of protein overexpression but reflects stabilization of the endogenous complex.

**Figure 6.**
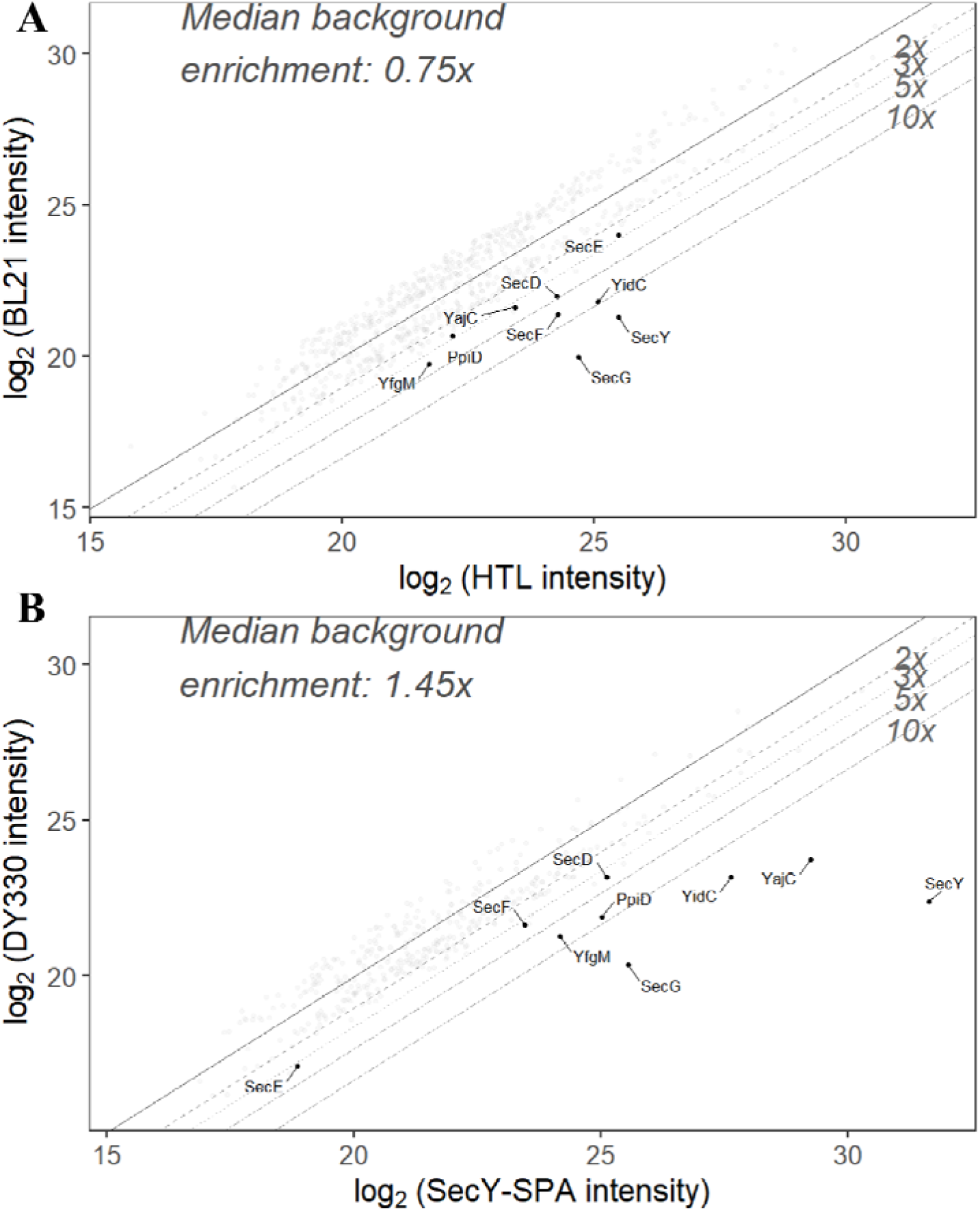
Affinity purification of the bacterial holo-translocon following PDET-1 extraction. (**A**) Scatter plot comparing protein group intensities in the affinity-purified holo-translocon and the corresponding BL21 control following PDET-1 extraction. The holo-translocon was overproduced from a plasmid in which SecE carries an N-terminal His tag, together with SecY, SecG, SecD, SecF, YajC, and YidC. Note that PpiD and YfgM were not encoded on the plasmid and were recovered as endogenous interaction partners. (**B**) Scatter plot comparing protein group intensities in the SecY-SPA affinity pulldown and the corresponding DY330 control following PDET-1 extraction. SecY was expressed from its native chromosomal locus. In both panels, the solid diagonal indicates equal abundance, and dashed lines indicate 2-, 3-, 5-, and 10-fold enrichment. Proteins are quantified via DIA-NN via DIA-MS.

Together, these findings establish that PDET-1 extends beyond detergent-free membrane proteomics by preserving membrane protein organization across multiple levels, from ligand binding and tissue-specific membrane proteomes to labile multi-subunit complexes, making it a versatile platform for functional membrane proteomics.

## Discussion

Quantitative membrane proteomics places unique demands on membrane extraction because extraction methods must preserve native membrane biology while remaining compatible with LC–MS/MS workflows. Although conventional detergents have been extensively optimized for membrane protein biochemistry and structural biology, peptide-mediated extraction has thus far been applied primarily to biochemical and structural studies of individual membrane proteins^16,18–20^. Here, we extend this approach by establishing Peptergents as a broadly applicable membrane extraction technology for LC–MS/MS-based membrane proteomics. Using the Peptergent PDET-1, we show that detergent-free extraction preserves ligand-responsive membrane proteins, stabilizes fragile endogenous membrane assemblies, and integrates seamlessly with membrane proteomic workflows.

Preservation of ligand-responsive membrane proteins provides one indication that native membrane organization is retained during extraction. ATP-vanadate stabilized MsbA only after PDET-1 extraction, whereas DDM-solubilized MsbA showed no detectable ligand-dependent thermal shift. Importantly, this observation extended from a purified bacterial transporter to the native mammalian membrane proteome, where the GPCR P2RY12 remained responsive to agonist binding following PDET-1 extraction but not DDM solubilization. Because thermal proteome profiling reports both ligand binding and the associated conformational response^6,28,29^, these findings indicate that peptide-mediated extraction preserves membrane proteins in states compatible with proteome-wide ligand discovery. Moreover, ligand responsiveness was retained following exchange into Peptidisc particles, demonstrating that the functional state established during extraction can be propagated into downstream membrane mimetics.

Preservation of native membrane organization also extended beyond individual proteins to higher-order membrane assemblies and tissue-specific membrane proteomes. The bacterial holo-translocon, a fragile nine-subunit complex that readily dissociates during detergent extraction^26,30^, remained intact following PDET-1 extraction from both overexpressing cells and chromosomally tagged strains. Likewise, peptide-mediated extraction preferentially recovered the CYP450 metabolic system together with its obligate redox partner POR and associated interaction network, while preserving tissue-specific membrane signatures in both liver and brain. Although these observations derive from distinct biological systems, they consistently indicate that peptide-mediated extraction preserves membrane organization across multiple biological scales, from ligand-responsive conformations and endogenous membrane assemblies to interaction networks and tissue-specific membrane proteomes.

Our quantitative proteomic analyses further demonstrate that peptide-mediated extraction and detergent solubilization optimize different aspects of membrane proteome recovery rather than simply representing weaker or stronger extraction methods. As expected, DDM produced deeper membrane proteome coverage, consistent with its greater membrane solubilization capacity. Nevertheless, both workflows recovered remarkably similar membrane protein composition, topology, functional classes, and tissue-specific biology, indicating that their principal differences lie in quantitative recovery rather than biological representation. DDM identified substantially more peptides, but these additional identifications were overwhelmingly of low abundance. In contrast, peptides shared between the two workflows consistently exhibited higher signal intensities following PDET-1 extraction, particularly among low- and moderately abundant species. Together with the preferential enrichment of CYP450 enzymes, POR, and tissue-specific membrane markers, these findings demonstrate that detergent-free extraction can preserve biologically coherent membrane proteomes while maintaining quantitative performance suitable for LC–MS/MS-based membrane proteomics.

The molecular basis of these recovery differences remains to be established. One contributing factor is likely the direct compatibility of peptide-mediated extraction with LC–MS/MS, which eliminates detergent removal and the associated sample losses inherent to conventional detergent workflows. Additional differences may arise from the distinct mechanisms by which peptides and detergents solubilize biological membranes. Unlike detergents, which assemble into relatively well-defined micelles, PDET-1 forms dynamic peptide–protein assemblies that can subsequently be exchanged into Peptidisc particles, suggesting a fundamentally different mode of membrane extraction. Similar dynamic behaviour, particularly lipid exchange, has been reported for other peptide-based nanodiscs^31^ and may contribute to the recovery biases observed with PDET-1. The preferential enrichment of endoplasmic reticulum proteins, including CYP450 enzymes and POR, may likewise reflect greater preservation of membrane environments that depend on specific phospholipid composition and organization^32,33^. Future studies will be required to determine how these physicochemical differences influence membrane protein recovery across diverse biological membranes.

In conclusion, our findings establish Peptergents as a broadly applicable membrane extraction technology that complements conventional detergent-based workflows for membrane proteomics. Rather than replacing detergents, Peptergents provide an alternative strategy for applications in which preserving ligand-responsive conformations, endogenous membrane assemblies, interaction networks, or tissue-specific membrane biology is particularly important. More broadly, our results highlight sample preparation as an important determinant of the biological information ultimately captured in membrane proteomic datasets. By preserving native membrane organization while remaining fully compatible with LC–MS/MS workflows, Peptergents expand the proteomics toolbox for studying membrane proteins and provide a robust technological foundation for future advances in membrane proteomics, interactomics, chemoproteomics, and structural biology.

## Materials and Methods

### Reagents

DDM was purchased from Anatrace (Cat no D310). Peptergent (purity 95%) and Peptidisc (purity 95%) were obtained from Peptidisc Biotech. Nickel nitrilotriacetic acid (Ni-NTA) chelating Sepharose resin was obtained from Qiagen (Cat no 30210) and ANTI-FLAG M2 Affinity Gel from Sigma (Cat no A2220). Silica beads (Cat no 440345) and the complete protease inhibitor cocktails were purchased from Sigma. Carboxylate-modified paramagnetic beads (Sera-Mag Speed-Beads (hydrophilic; Cat no 45152105050250) and Sera-Mag SpeedBeads (hydrophobic; Cat no 65152105050250) were purchased from GE Healthcare. 2-methylthio-ADP (2-MeSADP) was purchased from Cayman chemicals (Cat no 21230). Sodium orthovanadate (Cat no S454-50) as obtained from Fisher Scientific. Acetonitrile 99.9% (Cat no 271004), MS-grade trypsin (Cat no 90057) and DNAse (Cat no 10104159001) was purchased from Thermo Fisher Scientific. Octadecyl C18 Empore disks were purchased from 3M and Polygoprep 300-20 C18 powder was purchased from Macherey-Nagel. All other general reagents such as NaCl, Tris-base, ATP disodium trihydrate (Cat no: ATP007.5), DTT (DTT002.10), PMSF, iodoacetamide (IAA) (Cat no IOD500.5), urea, organic solvents, and acids were obtained from Bioshop.

### Bacterial strains and plasmids

Strain *E. coli* BL21(DE3) was obtained from our laboratory collection and used for recombinant protein expression. The *E. coli* sequential peptide affinity (SPA)-tagged strains used in this study were described previously^34^ and included the untagged parental strain DY330 and the SecY-SPA strain b3300. Plasmids encoding His□-tagged MsbA (pET28-His□-MsbA) and the HTL overexpression construct (pBAD22-HTL), carrying His□-SecE together with SecY, SecG, SecD, SecF, YajC, and YidC, were generated previously in our laboratory^26,35^.

### Protein expression and bacterial membrane preparation

His□-tagged MsbA and the His□-tagged HTL complex (pBAD22-HTL) were overproduced in *E. coli* BL21(DE3) grown at 37°C in 1 L of LB medium supplemented with antibiotic (kanamycin 25 μg/mL and ampicillin 50 μg/mL, respectively). Protein expression was induced with 0.5 mM IPTG (MsbA) or 0.2% arabinose during (HTL) exponential growth (OD□□□≈ 0.4 for MsbA and OD□□□≈ 1.0 for pBAD22-HTL). Similarly, BL21 parental control was grown at the same condition and time without antibiotic or induction. Cultures were incubated for an additional 3 h before cells were harvested by centrifugation (6,000 × g, 6 min). Cell pellets were resuspended in Buffer A (50 mM Tris-HCl, pH 7.8, 100 mM NaCl, 10% glycerol) supplemented with 1 mM phenylmethylsulfonyl fluoride (PMSF). Cells were disrupted using a microfluidizer (Microfluidics; three passes at 15,000 psi at 4°C). Unbroken cells and large debris were removed by centrifugation (6,000 × g, 6 min), and crude membranes were collected by ultracentrifugation (125,440g × g, 45 min, 4°C; Beckman Coulter Ti70 rotor). Membrane pellets were resuspended in Buffer A to a final protein concentration of 5 mg/mL and stored at −70°C until use.

### Tissue processing and preparation of crude membranes

All animal procedures were approved by the Animal Care Committee of the University of British Columbia and conducted in accordance with the Canadian Council on Animal Care guidelines.

Livers were collected from 12-week-old female C57BL/6 mice maintained under specific pathogen-free conditions and fed a standard chow diet. The authors acknowledge that the potential impact of mouse sex was not considered during study design. Following euthanasia, brains and livers were excised, rinsed with ice-cold phosphate-buffered saline (PBS) to remove residual blood, minced, and homogenized in ice-cold hypotonic lysis buffer (10 mM Tris-HCl, pH 7.4, 30 mM NaCl, 1 mM EDTA, 1× protease inhibitor cocktail, and 1 mM PMSF) using a tight-fitting metal Dounce homogenizer. Unless otherwise indicated, all subsequent steps were performed at 4°C or on ice. MgCl□(10 mM) and DNase I (50 μg/mL) were added, and the homogenate was incubated for 10 min before disruption by three passages through a French press (500 psi). Unbroken cells and nuclei were removed by centrifugation at 1,200 × g for 10 min, followed by centrifugation at 5,000 × g for 10 min to pellet mitochondria. Crude membranes were isolated by ultracentrifugation at 135,520g × g for 45 min (Beckman TLA110 rotor), resuspended in 100 μL TSG buffer (50 mM Tris-HCl, pH 7.9, 100 mM NaCl, 10% glycerol), and stored at −70°C until use.

### DDM and PDET-1 membrane solubilization

Crude membranes (∼1 mg protein) were resuspended in 0.5 mL of solubilization buffer containing 2 mg/mL Peptergent (PDET-1), 50 mM Tris-HCl (pH 7.8), 100 mM NaCl, 1 mM EDTA, 1 mM PMSF, and a protease inhibitor cocktail. Samples were incubated at 4°C for 60 min with gentle rotation. For detergent-based extraction, membranes were solubilized under identical conditions using 1% DDM. Insoluble material was removed by ultracentrifugation (135,520g × g, 15 min, 4°C), and the clarified supernatants were collected for downstream analyses.

### Affinity purification and Peptidisc reconstitution

Recombinant His□-tagged or FLAG-tagged membrane proteins were affinity purified from clarified membrane extracts prepared with PDET-1 or DDM. Clarified extracts were incubated with 100 μL of Ni-NTA agarose (His□-tagged proteins) or 50 μL anti-FLAG M2 affinity resin (FLAG-tagged proteins) for 1 h at 4°C with gentle mixing. After binding, the resins were washed three times with TS buffer (50 mM Tris-HCl, pH 7.8, 100 mM NaCl). For DDM-solubilized samples, the wash buffer was supplemented with 0.02% DDM to maintain protein solubility.

His□-tagged proteins were eluted with TS buffer containing 600 mM imidazole, whereas FLAG-tagged proteins were eluted with 100 mM glycine-HCl (pH 3.5; 100 µl). Glycine eluates were immediately neutralized by addition of 1 M Tris-HCl (pH 8.0; 10 µl). For Peptidisc reconstitution, Ni-NTA-bound His□-tagged proteins were incubated with (Fam-labeled) Peptidisc peptide (1 mg/mL) for 5 min at 4°C prior to elution. Excess peptide was removed by washing the resin once with TS buffer before eluting the protein with imidazole.

### Sodium orthovanadate preparation

Sodium orthovanadate (100 mM) was prepared by dissolving 36.8 mg sodium orthovanadate in 850 µL water at room temperature. The pH was adjusted to 10.0 by adding 6 M HCl dropwise (∼5 µL increments), at which point the solution turned yellow. The solution was boiled at 95°C for 10 min until it became colorless, cooled to room temperature, and the pH was re-adjusted to 10.0. This cycle of pH adjustment, boiling, and cooling was repeated two to three additional times until the solution remained colorless and the pH remained stable at 10.0 after boiling. The activated sodium orthovanadate solution was aliquoted and stored at −20°C.

### Thermal stability assay

Ligand-dependent thermal stabilization of purified MsbA was assessed using a thermal aggregation assay following membrane extraction with PDET-1, PDET-1 followed by Peptidisc reconstitution, or DDM. Purified MsbA was adjusted to approximately 0.5 mg/mL and incubated with 5 mM MgCl and 0.2 mM activated sodium orthovanadate. ATP was added to treatment samples to a final concentration of 2 mM, whereas control samples received an equivalent volume of water. Samples were incubated for 10 min at room temperature before thermal challenge. Two complementary thermal denaturation assays were performed. For kinetic measurements, samples were heated at 45 °C for 0–12 min. For thermal melting experiments, aliquots were heated for 3 min at temperatures ranging from 45 °C to 60 °C. Following heating, aggregated proteins were removed by ultracentrifugation (135,520g × g, 15 min, 4 °C; Beckman TLA100 rotor). Supernatants containing soluble MsbA were analyzed by SDS–PAGE. Ligand-dependent stabilization was assessed by comparing ATP/orthovanadate-treated samples with untreated controls.

### Thermal proteome profiling

Mouse liver crude membrane fractions were solubilized with DDM or PDET-1 as described above. For PDET-1 samples subsequently reconstituted into Peptidiscs, the Peptergent extract was exchanged into Peptidiscs by incubating the soluble fraction with a twofold excess (2 mg) of Peptidisc peptide for 10 min at 4°C. Solubilized extracts were then divided into ligand-treated and untreated samples. Ligand-treated samples received the selective P2RY12 agonist 2-methylthio-ADP (2-MeSADP) to a final concentration of 0.5 mM, whereas control samples received an equivalent volume of water. Following incubation for 10 min at room temperature, each condition was divided into two aliquots and heated independently at either 51 °C or 57 °C for 3 min. Precipitated proteins were removed by ultracentrifugation (135,520g × g, 15 min, 4 °C), and the soluble fractions were recovered for proteomic sample preparation. For comparison, thermal proteome profiling was also performed following conventional detergent extraction.

Crude membrane fractions were solubilized in 1% DDM under otherwise identical conditions. Ligand treatment, incubation, thermal challenge, ultracentrifugation, and downstream sample preparation were performed as described above.

### Native gel electrophoresis

Performed as previously described^36^, equal volumes of 4% and 12% acrylamide solutions were prepared in advance, and linear gradient gels were cast manually. TEMED and ammonium persulfate were added immediately before gradient mixing to initiate polymerization. Plastic combs were inserted, and the gels were allowed to polymerize for 30 min before storage at 4°C. For clear native PAGE, both the anode and cathode buffers consisted of buffer N (37 mM Tris-HCl, pH 8.8, 35 mM glycine).

### Proteomic sample preparation

Two sample preparation workflows were used depending on the membrane solubilization strategy. Because DDM is incompatible with LC–MS/MS, DDM-solubilized samples underwent SP4-based protein capture prior to digestion^37^, whereas Peptidisc-reconstituted samples were processed directly. For SP4, silica beads (9–13 µm diameter) were resuspended in water, washed once with 100% acetonitrile, rinsed twice with water, and resuspended at 50 mg/mL. Beads were pelleted by centrifugation (16,000 × *g*, 1 min) between washes. Protein samples (100 µg) were mixed with 1 mg silica beads, adjusted to 80% acetonitrile, incubated for 10 min at room temperature, and centrifuged (16,000 × *g*, 5 min) to immobilize proteins on the bead surface. The beads were washed three times with 80% ethanol without disturbing the pellet, then resuspended in 100 µL of 6 M urea. After centrifugation (16,000 × *g*, 1 min), the protein-containing supernatant was collected and processed according to the common digestion workflow described below. Peptidisc-reconstituted samples (100 µg protein) were denatured in 6 M urea for 30 min at room temperature. Both SP4-processed and Peptidisc-reconstituted samples were then subjected to the same digestion workflow. Briefly, proteins were reduced with 10 mM DTT for 1 h at room temperature, alkylated with 20 mM iodoacetamide (IAA) for 30 min in the dark, and quenched with 10 mM DTT for 30 min. Samples were diluted to 1 M urea with 50 mM ammonium bicarbonate (pH 8.0) and digested with MS-grade trypsin (Thermo Fisher Scientific; cat. no. 90057) at an enzyme-to-protein ratio of 1:100 for 24 h at 25°C with shaking. Digestion was terminated by acidification with formic acid to pH 3.0. Peptides were desalted using hand-packed C18 StageTips (Empore C18 disks), eluted with 40% acetonitrile containing 0.1% formic acid, and dried by vacuum centrifugation.

### LC–MS/MS Analysis

NanoLC connected to an Orbitrap Exploris 240 mass spectrometer (Thermo Fisher Scientific) was used for the analysis of all samples. The peptide separation was carried out using a Proxeon EASY nLC 1200 System (Thermo Fisher Scientific) fitted with a custom-made C18 column (15 cm x 150 μm ID) packed with HxSil C18 3 μm Resin 100 Å (Hamilton). A gradient of water/acetonitrile/0.1% formic acid was employed for chromatography. The samples were injected onto the column and run for 180 minutes at a flow rate of 0.60 μl/min. The peptide separation began with 1% acetonitrile, increasing to 3% in the first 4 minutes, followed by a linear gradient from 3% to 23% acetonitrile over 86 minutes, then another increase from 24% to 80% acetonitrile over 35 minutes, and finally a 35-minute wash at 80% acetonitrile, and then decreasing to 1% acetonitrile for 10 min and kept 1% acetonitrile for another 10 min. The eluted peptides were ionized using positive nanoelectrospray ionization (NSI) and directly introduced into the mass spectrometer with an ion source temperature set at 250°C and an ion spray voltage of 2.1 kV. In the case of data-dependent acquisition (DDA) mode, full-scan MS spectra (m/z 350–2000) were captured in Orbitrap Exploris 240 at a resolution of 120,000 (m/z 400). The automatic gain control was set to 1e6 for full FTMS scans and 5e4 for MS/MS scans. Ions with intensities above 1500 counts underwent fragmentation via NSI in the linear ion trap. The top 15 most intense ions with charge states of ≥2 were sequentially isolated and fragmented using normalized collision energy of 30%, activation Q of 0.250, and an activation time of 10 ms. Ions selected for MS/MS were excluded from further selection for 3 seconds. For EASY-nLC DIA, Orbitrap full MS scans were acquired from 375 to 1500 m/z at a resolution of 120,000 at m/z 400 with a normalized automated gain control (AGC) target of 300% and a maximum ion injection time set to Auto. For MS/MS scans, the HCD collision energy was set to 30%, the Orbitrap resolution to 15,000 at m/z 400, the normalized AGC target to 800%, the maximum injection time set to Auto, and the mass range to m/z 375 to 1500. For a theoretical cycle time of 3 s, 70 DIA windows of 19 m/z and an overlap of 0.5 m/z were used. The desired minimum points across the peak were set to 9, and the data type was centroid. The Orbitrap Exploris 240 mass spectrometer was operated using Thermo XCalibur software.

### DDA data processing with MaxQuant

Raw mass spectrometry data were processed using MaxQuant (v2.4.1.0)^38,39^. MS/MS spectra were searched with the Andromeda search engine against the UniProt Mus musculus reference proteome (UP000000589; December 2021; 55,086 entries) and the UniProt Escherichia coli reference proteome (UP000002032; July 2009; 4,156 entries). Initial precursor and fragment mass tolerances were both set to 20 ppm. Variable modifications included methionine oxidation, asparagine/glutamine deamidation, and protein N-terminal acetylation, while carbamidomethylation of cysteine was specified as a fixed modification. A maximum of two missed tryptic cleavages was allowed, with a minimum peptide length of six amino acids. Protein identification was performed using a target-decoy strategy based on an automatically generated reverse database, with the false discovery rate (FDR) controlled at 1% for both peptide-spectrum matches and protein identifications. Proteins identified exclusively by shared peptides were reported as a single protein group. Label-free quantification (LFQ) was performed using the MaxLFQ algorithm implemented in MaxQuant.

### DIA data processing with DIA-NN

Data-independent acquisition (DIA) datasets were processed using DIA-NN^40^ v1.9.1 in library-free mode with contaminant identification enabled. A predicted spectral library was generated directly from the UniProt Escherichia coli reference proteome (UP000000625; December 2025 release) using DIA-NN’s integrated deep learning algorithms to predict fragment ion spectra, retention times (RTs), and ion mobilities (IMs). Protein digestion was specified using Trypsin/P with a maximum of one missed cleavage. Carbamidomethylation of cysteine was set as a fixed modification, and no variable modifications were permitted. N-terminal methionine excision was enabled. Peptide lengths were restricted to 7–30 amino acids, precursor charges to 1–4, precursor m/z values to 300–1,800, and fragment ion m/z values to 200–1,800. Protein inference was performed at the gene level using heuristic inference with shared spectra excluded. Precursor-level false discovery rates (FDRs) were controlled at 1%, the integrated neural network classifier was run in single-pass mode, and the log level was set to 1. Quantification was performed using the QuantUMS high-precision algorithm with retention time-dependent cross-run normalization enabled. All DIA datasets were processed simultaneously in a single DIA-NN analysis to maximize identification consistency and quantitative accuracy across all biological samples.

### Figure Generation, Data Analysis, and Code Availability

The graphical abstract was created using BioRender. Figures 1C, 2C, and Supplemental Figure 3B were generated using GraphPad Prism (v.11.0.2). Figures 3–6, Table 1, Supplemental Figures 4–11, and Supplemental Table 1 were generated in R (v.4.5.1) using RStudio (v.2026.7.0.139).

All volcano plot significance testing was performed using two-tailed t-tests in Perseus^41^ (v.3); all other statistical analyses were performed in R. R packages used include tidyverse^42^, ggsignif, ggrepel, patchwork, cowplot, gt, webshot2, boot, clusterProfiler^43^, BiocManager, org.Mm.eg.db^44^, AnnotationDbi, GO.db, homologene, TissueEnrich, GSEABase, SummarizedExperiment, readxl, and STRINGdb. Transmembrane segment prediction was performed using Phobius^45^ and GO terms from UniProt^46^. Densitometry analysis in Figures 1C, 2C and Supplemental Figure 3B were performed with ImageJ (v 1.54t)^47^.

## CRediT authorship contribution statement

Conceptualization: F.A., F.D.v.H.

Methodology: F.A.

Validation: F.A., A.B.

Formal Analysis: F.A., A.B.

Investigation: F.A. H.A.

Writing - Original Draft: F.A., A.B.

Writing - Review & Editing: A.B. F.D.v.H.

Visualization: F.A., A.B.

Supervision: F.D.v.H., M.B.

Project Administration: F.A., F.D.v.H., M.B.

Funding Acquisition: A.B., F.D.v.H., M.B.

## Acknowledgments

Work in the Duong lab was supported by an NSERC Discovery grant. Work in the Babu lab was supported by the Canada Foundation for Innovation and CIHR Foundation grant FDN-154318. A. B. is supported by a UBC 4-year fellowship, an Amplify Doctoral Award from Triangle (Training a new generation of researchers in gastroenterology and liver), and a Mast3 (Mass Spectrometry Team Training and Transition) scholarship.

## Notes

The mass spectrometry proteomics data have been deposited in the ProteomeXchange Consortium via the PRIDE partner repository. Accession and Token available upon demand to the corresponding author(s)

## SUPPLEMENTAL FIGURES

**Supplemental Figure 1.**
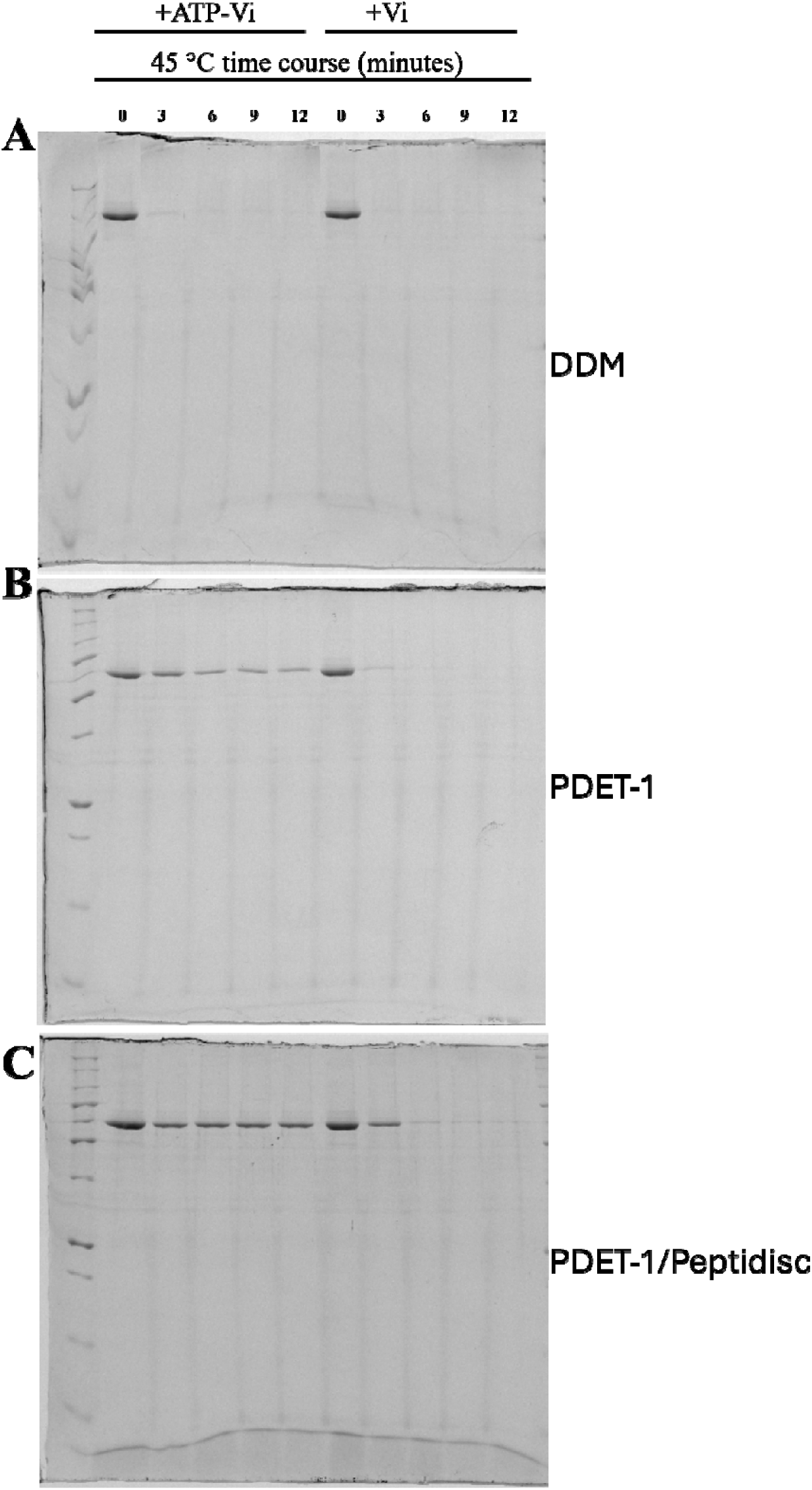
Uncropped SDS–PAGE gels for all samples from the MsbA ATP–vanadate time-course experiment shown in Figure 2.

**Supplemental Figure 2.**
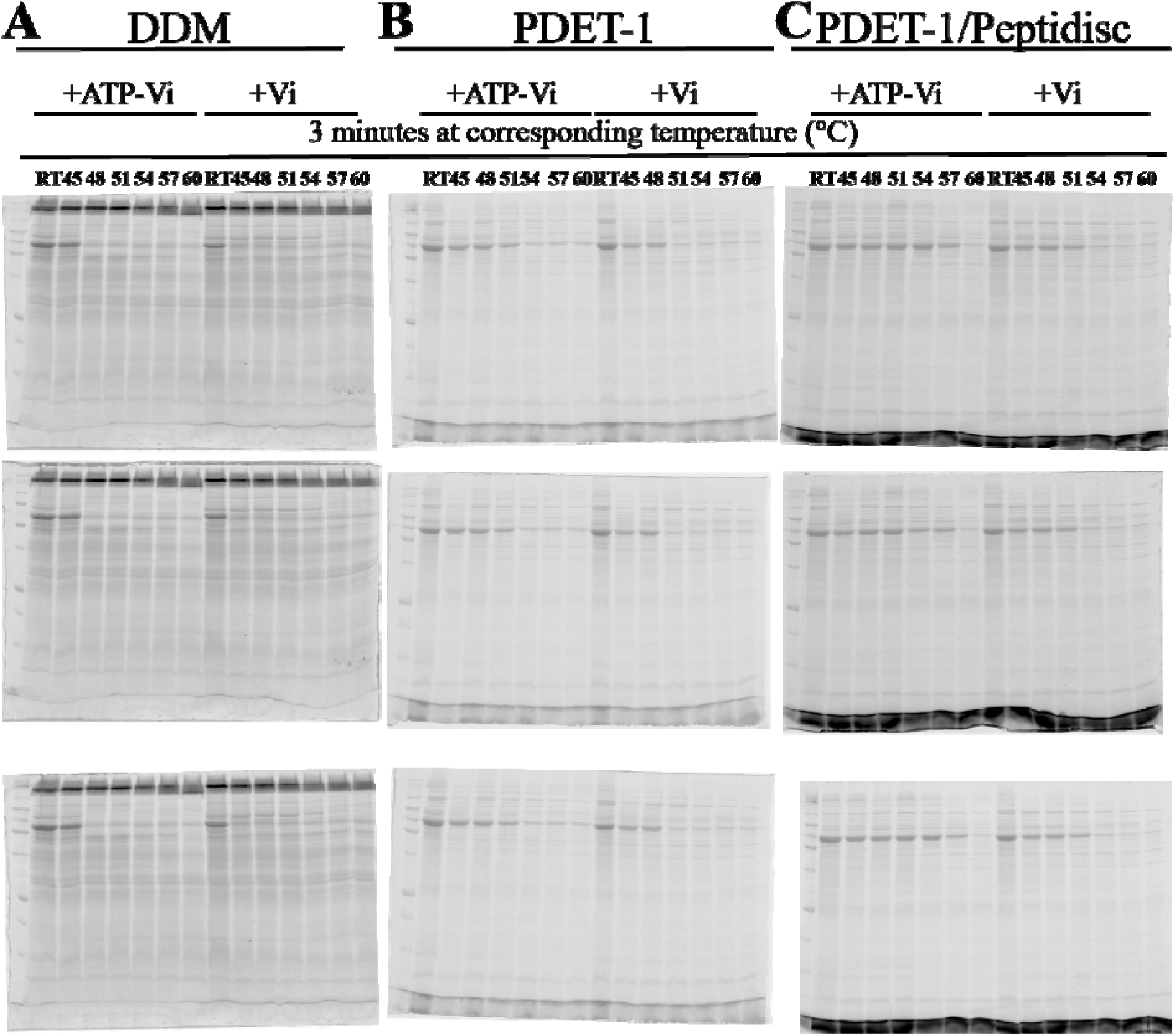
Uncropped SDS–PAGE gels for all samples from the MsbA ATP–vanadate time-course experiment shown in Figure 2.

**Supplemental Figure 3.**
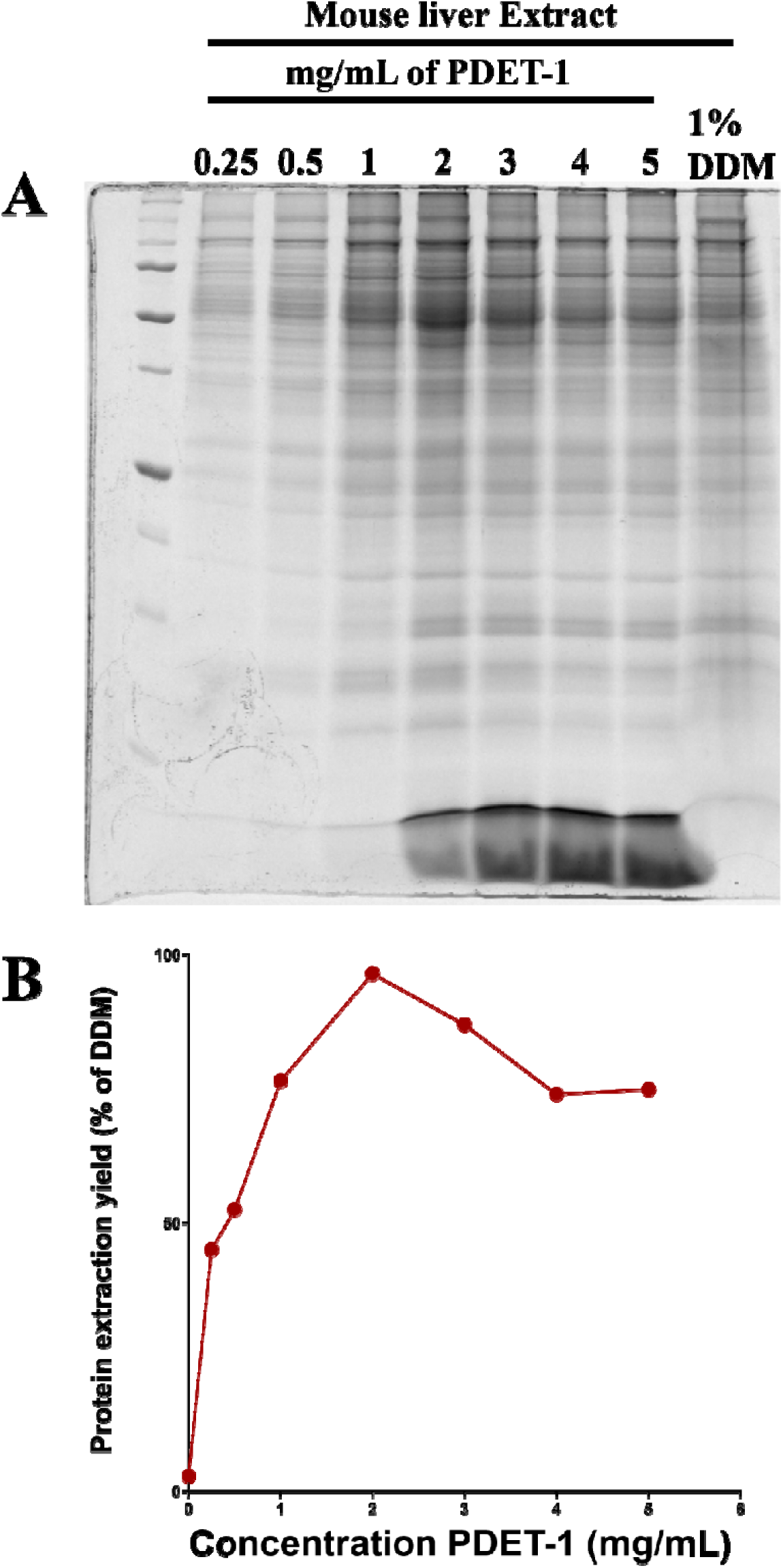
Optimization of PDET-1 extraction from mouse liver membranes. (**A**) Coomassie-stained SDS-PAGE gel showing mouse liver membrane proteins extracted with increasing concentrations of PDET-1 (0.25, 0.5, 1, 2, 4, and 8 mg/mL). DDM extraction (1%) is shown as a reference. (**B**) Protein extraction yield, measured by densitometry scanning of the gel with ImageJ from (A) and expressed as a percentage of the yield obtained with DDM.

**Supplemental Figure 4.**
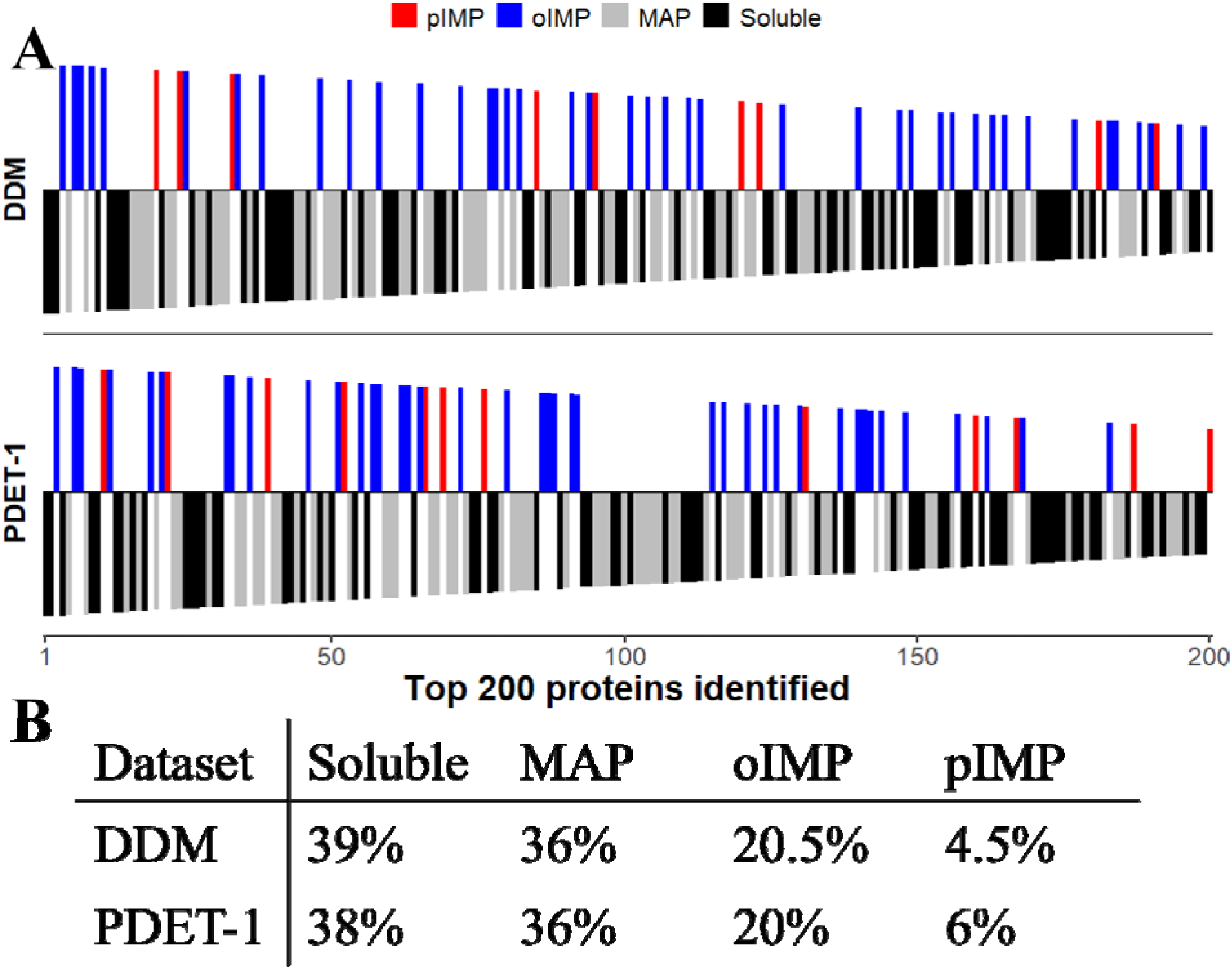
Subcellular localization of the 200 most abundant mouse liver proteins recovered by DDM and PDET-1. **(A)** Barcode plots showing the subcellular localization of the 200 most abundant proteins (ranked by mean intensity across biological replicates) recovered from mouse liver membranes following extraction with DDM (top) or PDET-1 (bottom). Each bar represents a single protein and is colored according to localization category. IMPs are plotted above the horizontal axis, whereas MAP and soluble proteins are plotted below. (**B**) Percentage of proteins assigned to each localization category of the top 200 most abundant proteins.

**Supplemental Figure 5.**
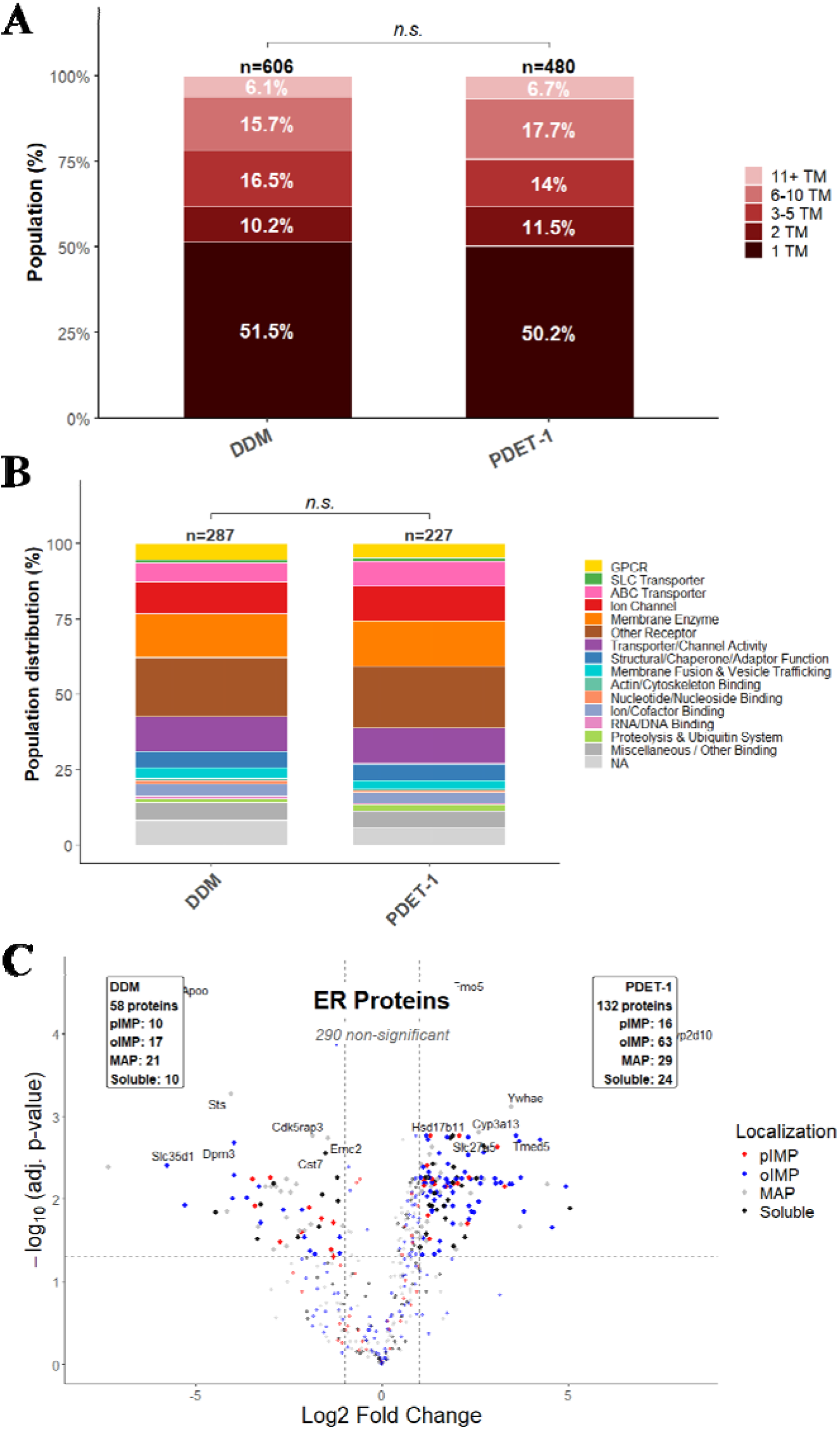
PDET-1 and DDM recover membrane proteins with similar transmembrane topologies and plasma membrane functional classes but differ in recovery of ER proteins. **(A)** Distribution of transmembrane (TM) helix numbers among integral membrane proteins recovered from mouse liver membranes following extraction with DDM (n = 606) or PDET-1 (n = 480). Proteins were grouped into five TM classes (1, 2, 3–5, 6–10, and ≥11 TM helices). The TM distributions did not differ significantly between extraction methods (χ² test, n.s.). Post-hoc Fisher’s exact tests within each TM bin (BH-adjusted) likewise showed no significant differences (1 TM: p.adj = 0.714; 2 TM: p.adj = 0.714; 3–5 TM: p.adj = 0.714; 6–10 TM: p.adj = 0.714; 11+ TM: p.adj = 0.714). **(B)** Functional classification of plasma membrane integral membrane proteins (pIMPs) recovered by DDM (n = 262) or PDET-1 (n = 227). Proteins were assigned to functional classes based on Gene Ontology molecular function and subcellular localization annotations. The functional distributions were not significantly different between extraction methods (χ² = 2.64, df = 15, p = 0.9998; Fisher’s exact test with simulated p-value based on 10,000 Monte Carlo replicates, p = 0.9996). **(C)** Volcano plot of endoplasmic reticulum (ER)-annotated proteins (n = 480; smooth ER and ER GO cellular component terms, merged) comparing PDET-1 and DDM extraction (log□fold change versus −log adjusted p-value).

**Supplemental Figure 6.**
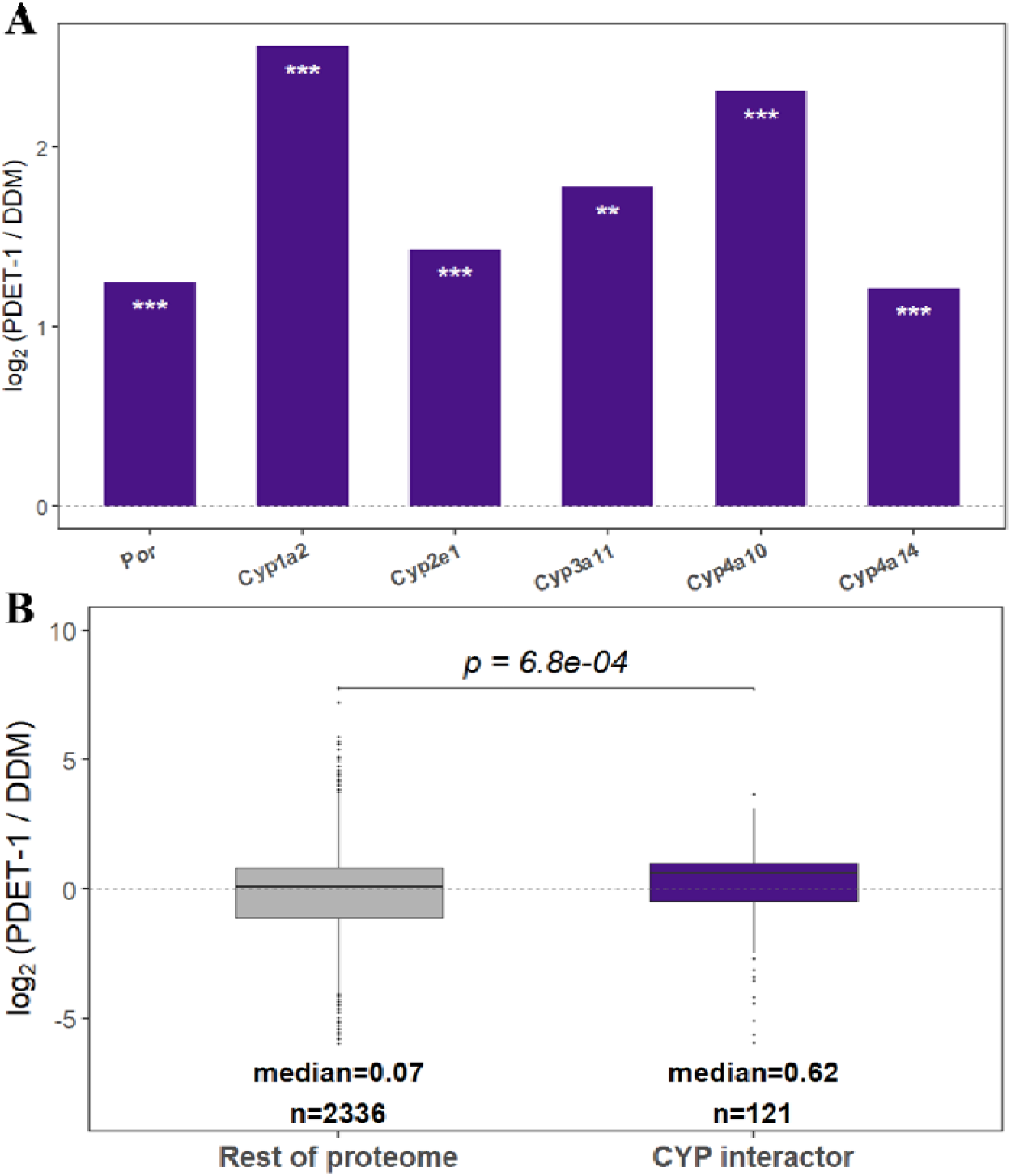
PDET-1 preferentially enriches cytochrome P450 enzymes and their known protein interactors relative to DDM. **(A)** Bar graph showing the log fold change (PDET-1/DDM) for individual cytochrome P450 enzymes (Cyp1a2, Cyp2e1, Cyp3a11, Cyp4a10, and Cyp4a14) and their obligate redox partner Por, all of which are significantly enriched by PDET-1 (Cyp1a2: log FC = 2.56, p = 6.31 × 10□□; Cyp4a10: log FC = 2.31, p = 2.04 × 10□□; Cyp3a11: log FC = 1.78, p = 1.15 × 10□³; Cyp2e1: log FC = 1.43, p = 2.95 × 10□v; Por: log FC = 1.25, p = 2.19 × 10□□; Cyp4a14: log FC = 1.21, p = 3.24 × 10). Positive values indicate enrichment by PDET-1. Asterisks denote statistical significance (**, p < 0.01; \*\**, p < 0.001).* **(B)** Box plot comparing the distribution of log fold changes (PDET-1/DDM) for proteins annotated as cytochrome P450 interactors (n = 121; high-confidence STRING interactions, combined score ≥700) and all other quantified proteins (n = 2,336). CYP interactors showed a higher median log FC (0.62) than the rest of the proteome (0.07). Cytochrome P450 interactors exhibit significantly greater enrichment by PDET-1 than the remainder of the proteome (Wilcoxon rank-sum test, W = 167,169, p = 6.8 × 10□□).

**Supplemental Figure 7.**
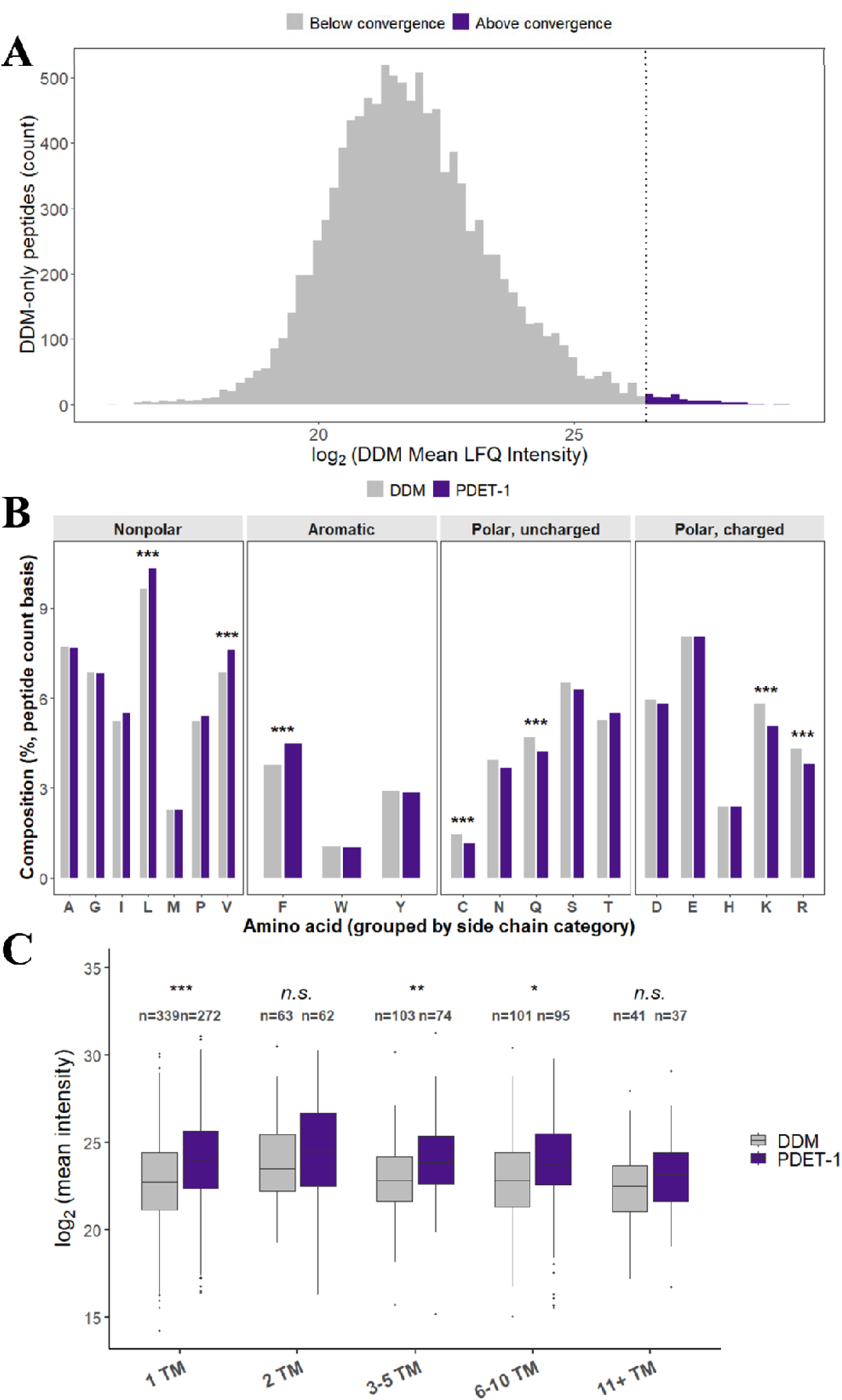
DDM-specific peptides are predominantly low abundance, whereas PDET-1 preferentially recovers more hydrophobic peptides with higher signal intensities. **(A)** Histogram showing the distribution of mean log □LFQ intensities for peptides identified exclusively by DDM (n = 11,047). The dashed vertical line denotes the empirical LFQ convergence threshold (∼8.8 × 10). Peptides below the threshold are shown in gray and those above in purple. Of the DDM-specific peptides, 99.1% (10,951 of 11,047) fell below the convergence threshold, indicating that DDM’s greater peptide coverage is driven almost entirely by low-abundance peptides. **(B)** Relative amino acid composition of peptides identified exclusively by DDM or PDET-1, grouped by side-chain class (nonpolar, aromatic, polar uncharged, and charged). Overall amino acid composition differed significantly between DDM-only and PDET-1-only peptides (χ² = 165.8, df = 19, p < 2.2 × 10 ¹; Cramér’s V = 0.029, indicating a small effect size). Statistical significance for individual amino acids was assessed using a two-proportion z-test with Benjamini–Hochberg correction across all 20 amino acids (adjusted p < 0.05; **, adjusted p < 0.01; **, adjusted p < 0.001). Significant differences were observed for phenylalanine (F, p.adj = 1.6 × 10□□), lysine (K, p.adj = 9.3 × 10□v), valine (V, p.adj = 4.5 × 10□□), cysteine (C, p.adj = 6.0 × 10 □□), arginine (R, p.adj = 1.5 × 10□□), glutamine (Q, p.adj = 3.9 × 10□□), and leucine (L, p.adj = 6.3 × 10□□); all other amino acids were not significant after correction. **(C)** Distribution of mean log□ peptide intensities for peptides mapping to transmembrane integral membrane proteins, grouped according to the number of transmembrane (TM) helices (1, 2, 3–5, 6–10, and ≥11). The number of peptides (n) in each group is indicated above the corresponding box plot. Statistical significance was assessed using Wilcoxon rank-sum tests with Benjamini–Hochberg correction across TM bins (1 TM: p.adj = 3.5 × 10□□; 2 TM: p.adj = 0.278; 3–5 TM: p.adj = 7.8 × 10 ³; 6–10 TM: p.adj = 1.3 × 10□²; 11+ TM: p.adj = 0.132; n.s., not significant; *, adjusted p < 0.05; **, adjusted p < 0.01;**, adjusted p < 0.001).

**Supplemental Figure 8.**
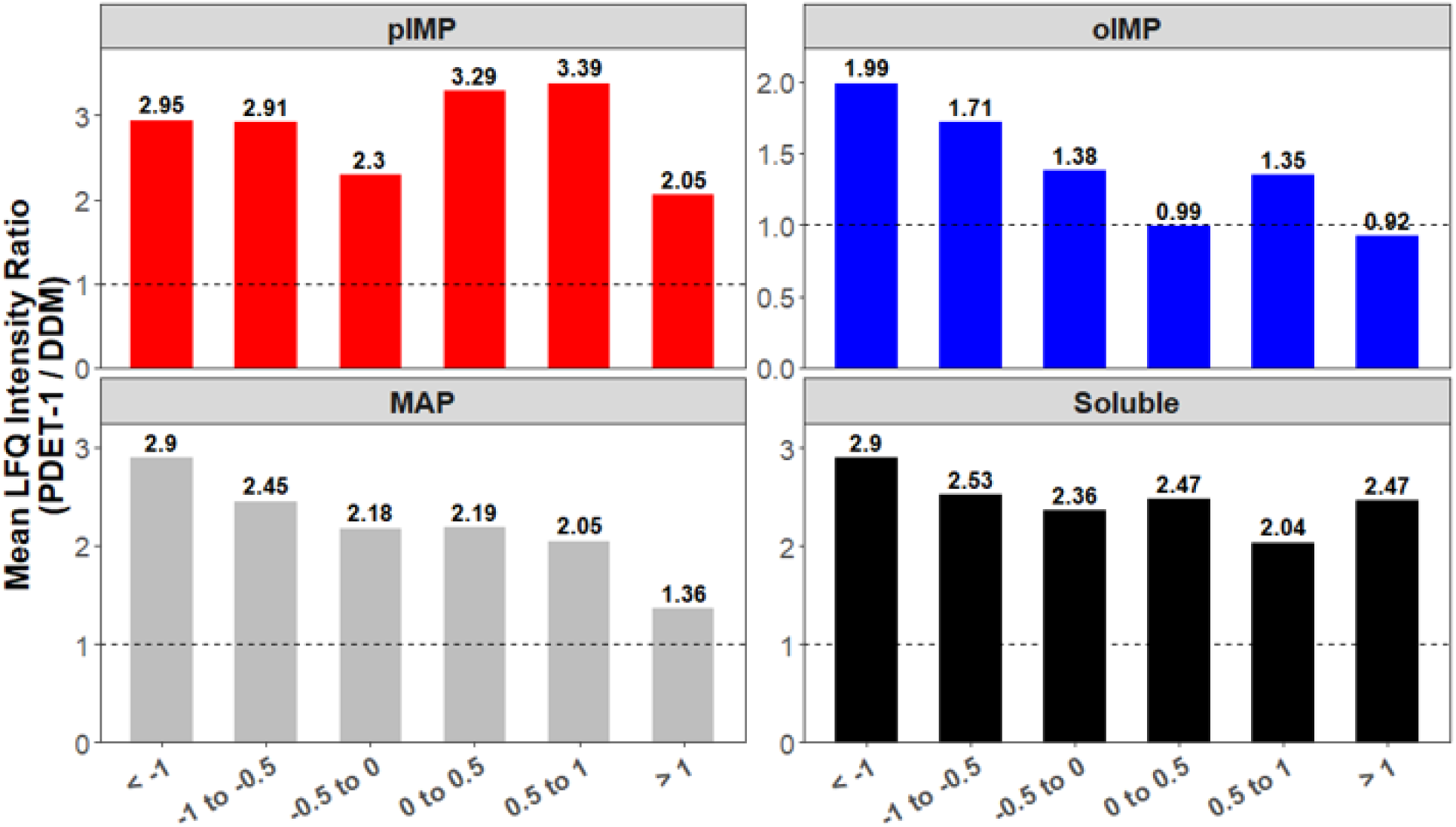
PDET-1 enhances peptide intensity across the hydrophobicity spectrum, with the greatest relative gains in pIMPs and the most modest gains in oIMPs. Bar graphs showing the mean LFQ intensity ratio (PDET-1/DDM) across peptide hydrophobicity bins defined by GRAVY (grand average of hydropathicity) score (< −1, −1 to −0.5, −0.5 to 0, 0 to 0.5, 0.5 to 1, and >1), stratified by protein localization category. The dashed horizontal line indicates an intensity ratio of 1 (equal mean LFQ intensity between extraction methods), and the corresponding ratio is shown above each bar. Peptides detected in at least two of three biological replicates were included in the analysis and values above 1 indicate greater mean peptide intensity in PDET-1 relative to DDM. PDET-1 shows enhanced intensity relative to DDM across nearly all GRAVY bins and localization categories, with pIMP showing the largest relative gains (ratios ranging from 2.05 to 3.39 across bins). MAP and soluble proteins show intermediate, broadly similar gains (ratios of 1.36–2.90 and 2.04–2.90, respectively), while organellar integral membrane proteins (oIMP) show the most modest gains and, at the highest GRAVY bins, near-equivalent or slightly DDM-favoring ratios (0.988 at 0 to 0.5; 0.925 at > 1).

**Supplemental Table 1.** Proteome coverage and subcellular localization of mouse brain membrane proteins solubilized with DDM and PDET-1. Total protein counts and localization distribution for mouse brain membrane proteins extracted with DDM or PDET-1 and analyzed by MaxQuant.

| MS sample<br>(mean ± SD) | Total proteins<br>(TPs) | TPs | | Integral membrane proteins<br>(tIMPs) | IMPs | |
| --- | --- | --- | --- | --- | --- | --- |
|  |  | Soluble proteins<br>(SPs) | Membrane associated proteins<br>(MAPs) |  | pIMPs | oIMPs |
| mBrain DDM | 2984 ± 19 | 1278 ± 8<br>(42.8%) | 1706 ± 11<br>(57.2%) | 739 ± 2<br>(24.8%) | 389 ± 6<br>(13.0%) | 350 ± 4<br>(11.7%) |
| mBrain PDET-1 | 2071 ± 72 | 822 ± 42<br>(39.7%) | 1250 ± 30<br>(60.3%) | 542 ± 15<br>(26.2%) | 289 ± 7<br>(14.0%) | 253 ± 9<br>(12.2%) |

**Supplemental Figure 9.**
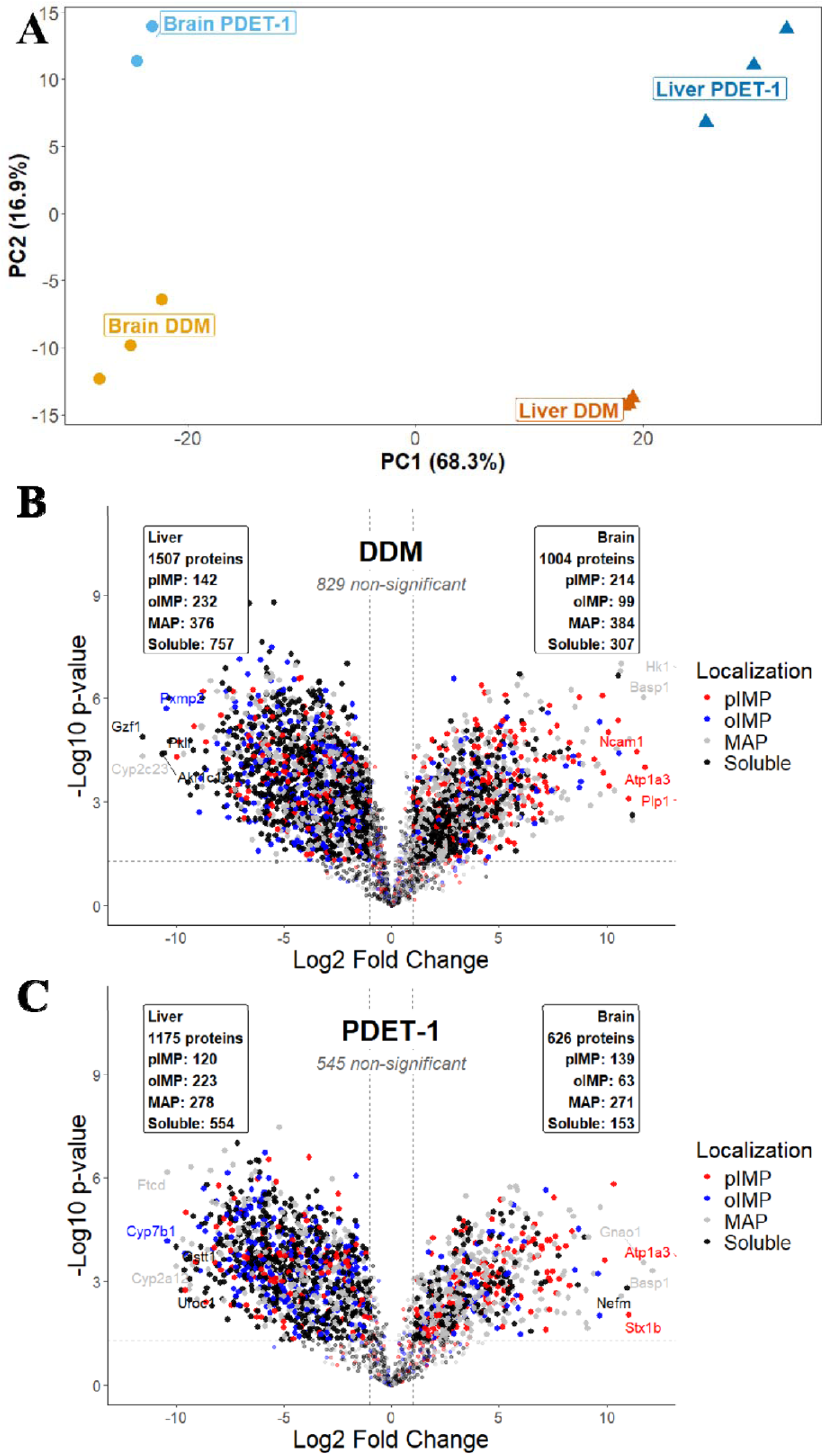
Tissue identity is the primary determinant of membrane proteome composition, with extraction method contributing secondary variation. **(A)** Principal component analysis (PCA) of mouse brain and liver membrane proteomes extracted with DDM or PDET-1. Samples are colored by extraction method and tissue. PC1 (68.3% of the variance) separates samples by tissue, whereas PC2 (16.9%) distinguishes samples by extraction method. **(B)** Volcano plot comparing liver and brain membrane proteomes extracted with PDET-1 (log□fold change versus −log□□*p*-value). **(C)** Volcano plot comparing log_2_ fold change between liver and brain membrane proteomes in the DDM dataset, formatted as in (B).

**Supplemental Figure 10.**
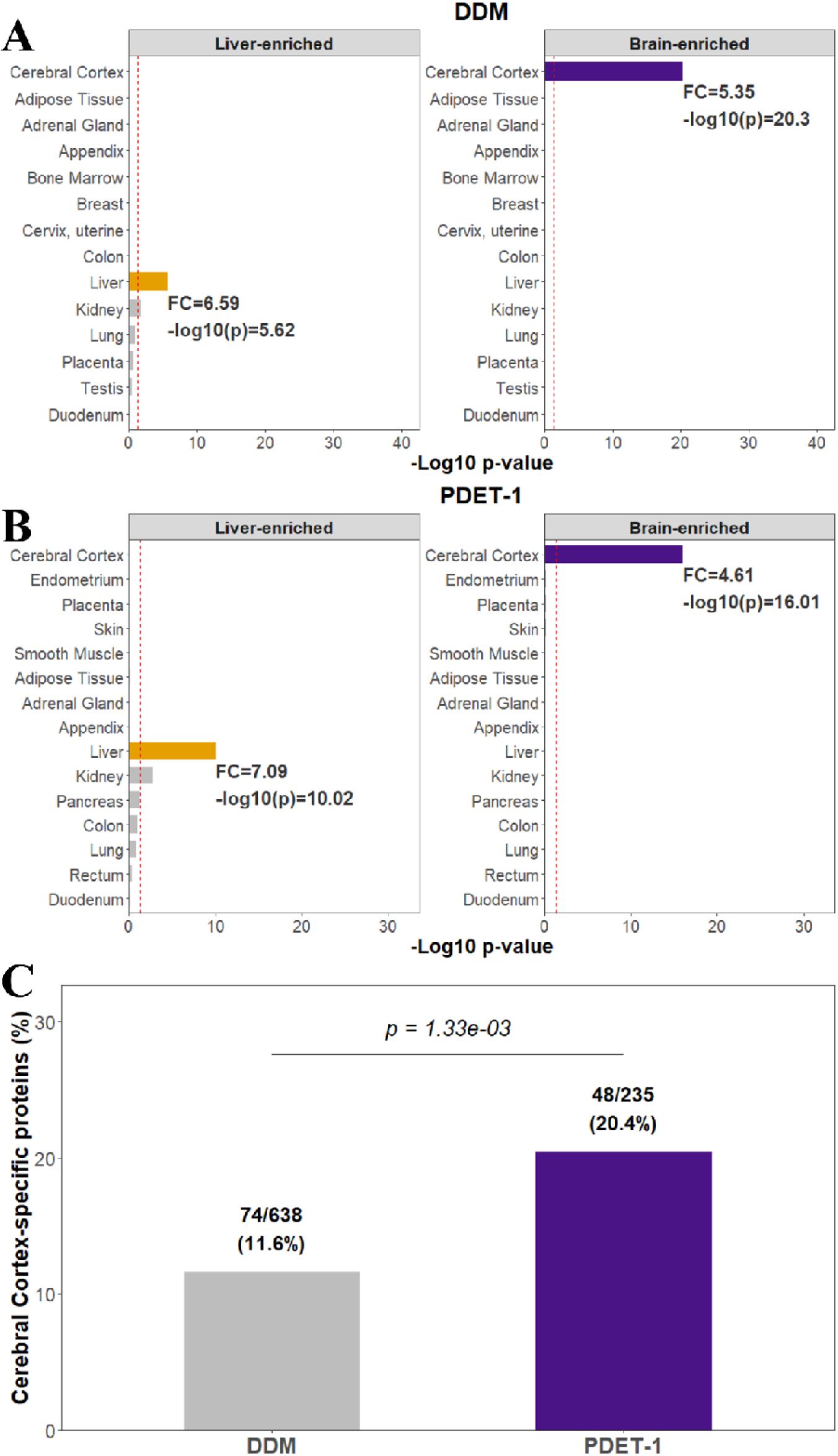
PDET-1 enhances enrichment of tissue-specific markers in mouse liver and brain membrane proteomes. **(A)** Tissue enrichment analysis (TissueEnrich; GTEx reference) of proteins enriched in liver (left) or brain (right) following DDM extraction. Bars represent the enrichment significance (–log□□p-value, hypergeometric test) for each reference tissue; the dashed vertical line indicates the significance threshold (p = 0.05). Liver-enriched proteins showed the strongest enrichment for the liver (fold change = 6.59, −log10(p) = 5.62), whereas brain-enriched proteins showed the strongest enrichment for the cerebral cortex (fold change = 5.35, −log10(p) = 20.30). **(B)** Tissue enrichment analysis of proteins enriched following PDET-1 extraction, formatted as in (A). Liver and cerebral cortex again showed the strongest enrichment for liver- and brain-enriched proteins (FC = 7.09, −log10(p) = 10.02 and FC = 4.61, −log10(p) = 16.01, respectively). **(C)** Percentage of DDM- or PDET-1-enriched brain proteins (identified by comparing DDM and PDET-1 extraction directly within brain membranesr) annotated as cerebral cortex-specific markers by TissueEnrich. Values above the bars indicate the number of cerebral cortex markers relative to the total number of significantly enriched proteins for each extraction method. PDET-1 recovered a significantly greater proportion of cerebral cortex-specific markers than DDM (Fisher’s exact test, OR = 1.95, p = 0.0013).

**Supplemental Figure 11.**
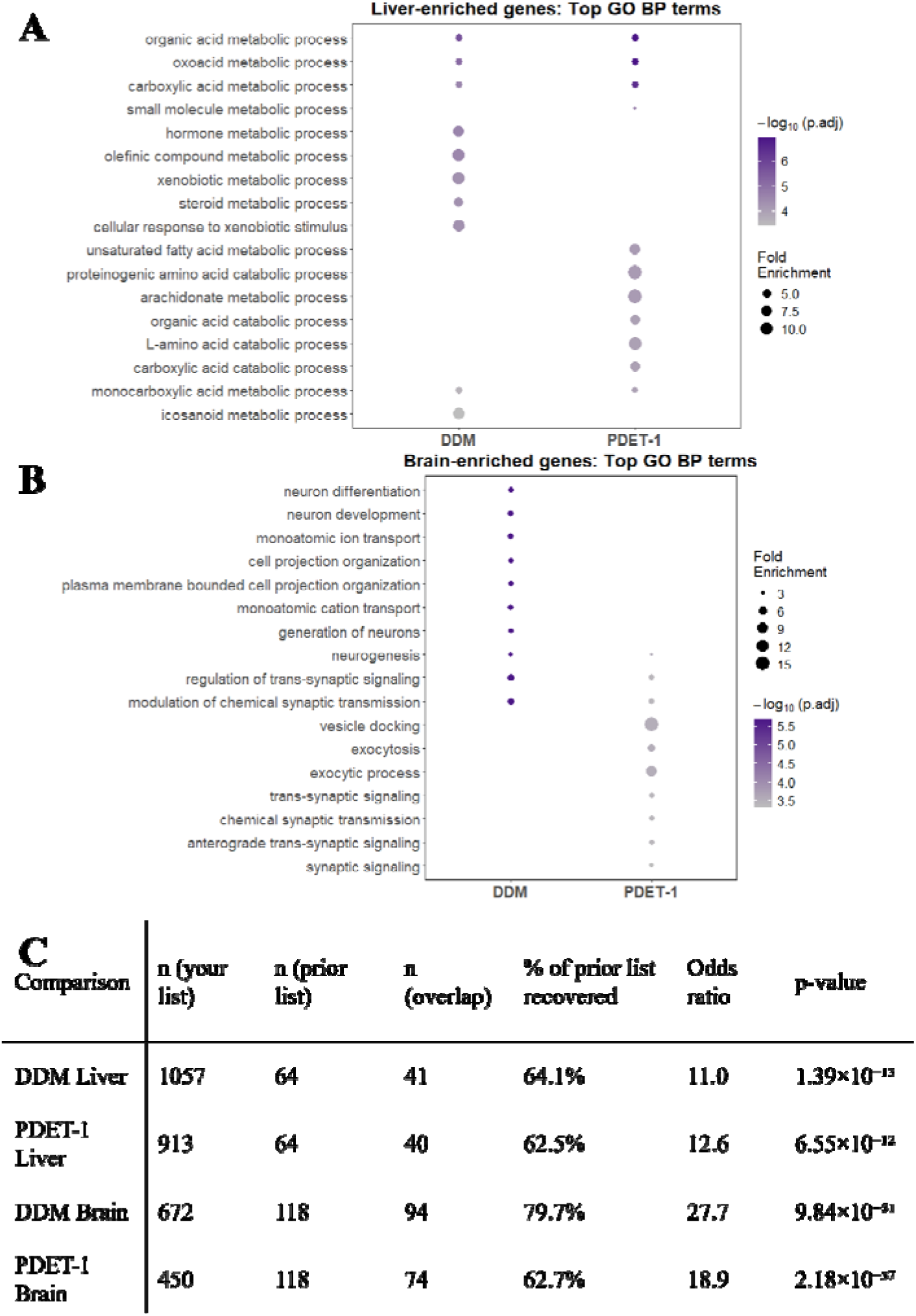
Tissue-enriched proteins recovered by DDM and PDET-1 exhibit expected biological functions and significantly overlap with previously published tissue-specific plasma membrane proteins. **(A)** Dot plot showing the top enriched Gene Ontology Biological Process (GO BP) terms for brain-enriched proteins recovered by DDM or PDET-1. Dot size represents fold enrichment and color indicates enrichment significance (–log□□ adjusted p-value, hypergeometric test with benjamini hochberg correction within each gene set comparison). Both extraction methods show enrichment for processes associated with neuronal development, cell projection organization, ion transport, and synaptic signaling. **(B)** GO BP enrichment analysis of liver-enriched proteins recovered by DDM or PDET-1, formatted as in (A). Both datasets show enrichment for metabolic processes characteristic of the liver, including organic acid, carboxylic acid, xenobiotic, and steroid metabolism. **(C)** Overlap between tissue-enriched protein sets identified in this study and a previously published reference set of tissue-enriched plasma membrane integral membrane proteins (pIMPs). For each comparison, the total number of proteins, the number of overlapping proteins, the percentage of the reference set recovered, the odds ratio, and Fisher’s exact test p-value are shown. All four datasets exhibit highly significant overlap with the published tissue-specific pIMP reference sets.

## REFERENCES

1. Uhlén, M. et al. Tissue-based map of the human proteome. Science 347, 1260419 (2015).

2. Van ‘T Klooster, J. S., et al. Periprotein lipidomes of Saccharomyces cerevisiae provide a flexible environment for conformational changes of membrane proteins. eLife 9, e57003 (2020).

3. Tan, S., Tan, H. T. & Chung, M. C. M. Membrane proteins and membrane proteomics. PROTEOMICS 8, 3924–3932 (2008).

4. Brough, Z., Zhao, Z. & Duong Van Hoa, F. From bottom-up to cell surface proteomics: detergents or no detergents, that is the question. Biochem. Soc. Trans. 52, 1253–1263 (2024).

5. Lambos, J. C., Bhattacharya, A., Al-Seragi, M. & Duong Van Hoa, F. Expanding the Reach of Membrane Protein–Ligand Interaction Studies Through the Integration of Mass Spectrometry and Membrane Mimetics. PROTEOMICS 25, 25–40 (2025).

6. Jandu, R. S. et al. Membrane-mimetic thermal proteome profiling (MM-TPP) toward mapping membrane protein–ligand dynamic interactions. eLife 14, RP104549 (2025).

7. Antony, F. et al. Comparative Evaluation of Solid-phase and Membrane Mimetic Strategies in Membrane Proteome Coverage and Disease-State Analysis. Mol. Cell. Proteomics 25, 101496 (2026).

8. Tribet, C., Audebert, R. & Popot, J.-L. Amphipols: Polymers that keep membrane proteins soluble in aqueous solutions. Proc. Natl. Acad. Sci. 93, 15047–15050 (1996).

9. Brown, C. et al. A proteome-wide quantitative platform for nanoscale spatially resolved extraction of membrane proteins into native nanodiscs. Nat. Methods 22, 412–421 (2025).

10. Zhang, S. et al. One-step construction of circularized nanodiscs using SpyCatcher-SpyTag. Nat. Commun. 12, 5451 (2021).

11. Mostafavi, S., Custódio, T. F., Jungnickel, K. E. J. & Löw, C. Salipro technology in membrane protein research. Curr. Opin. Struct. Biol. 93, 103050 (2025).

12. Zhao, Z. et al. A Peptidisc-Based Survey of the Plasma Membrane Proteome of a Mammalian Cell. Mol. Cell. Proteomics 22, 100588 (2023).

13. Carlson, M. L. et al. The Peptidisc, a simple method for stabilizing membrane proteins in detergent-free solution. eLife 7, e34085 (2018).

14. Bhattacharya, A. et al. Membrane Proteome Remodeling in Female APP/PS1 Mice Following M1 Muscarinic Receptor Modulation Revealed by Peptidisc-Enabled DIA-MS. J. Proteome Res. 25, 3136–3148 (2026).

15. Antony, F. et al. Sensitive Profiling of Mouse Liver Membrane Proteome Dysregulation Following a High-Fat and Alcohol Diet Treatment. PROTEOMICS 24, e202300599 (2024).

16. Zang, J. et al. Unveiling Eukaryotic Membrane Proteins in High Resolution Using Peptide Solubilization. J. Mol. Biol. 437, 169467 (2025).

17. Antony, F., Bhattacharya, A. & Duong Van Hoa, F. PEPTERGENT: A Peptide-Based Reagent for Detergent-Free Extraction of Membrane Proteins and Purification of Membrane Proteomes. BIO-Protoc. 16, (2026).

18. Yeh, J. I., Du, S., Tortajada, A., Paulo, J. & Zhang, S. Peptergents: Peptide Detergents That Improve Stability and Functionality of a Membrane Protein, Glycerol-3-phosphate Dehydrogenase. Biochemistry 44, 16912–16919 (2005).

19. Zhao, X. et al. Designer short peptide surfactants stabilize G protein-coupled receptor bovine rhodopsin. Proc. Natl. Acad. Sci. 103, 17707–17712 (2006).

20. Kiley, P. et al. Self-Assembling Peptide Detergents Stabilize Isolated Photosystem Ion a Dry Surface for an Extended Time. PLoS Biol. 3, e230 (2005).

21. Zhang, J. et al. Agonist-bound structure of the human P2Y12 receptor. Nature 509, 119–122 (2014).

22. Riddick, D. S. et al. NADPH–Cytochrome P450 Oxidoreductase: Roles in Physiology, Pharmacology, and Toxicology. Drug Metab. Dispos. 41, 12–23 (2013).

23. Szklarczyk, D. et al. The STRING database in 2025: protein networks with directionality of regulation. Nucleic Acids Res. 53, D730–D737 (2025).

24. Jain, A. & Tuteja, G. TissueEnrich: Tissue-specific gene enrichment analysis. Bioinformatics 35, 1966–1967 (2019).

25. Antony, F., Brough, Z., Zhao, Z. & Duong Van Hoa, F. Capture of the Mouse Organ Membrane Proteome Specificity in Peptidisc Libraries. J. Proteome Res. 23, 857–867 (2024).

26. Young, J. W. et al. Development of a Method Combining Peptidiscs and Proteomics to Identify, Stabilize, and Purify a Detergent-Sensitive Membrane Protein Assembly. J. Proteome Res. 21, 1748–1758 (2022).

27. Troman, L. A. & Collinson, I. Pushing the Envelope: The Mysterious Journey Through the Bacterial Secretory Machinery, and Beyond. Front. Microbiol. 12, 782900 (2021).

28. Savitski, M. M. et al. Tracking cancer drugs in living cells by thermal profiling of the proteome. Science 346, 1255784 (2014).

29. Kalxdorf, M. et al. Cell surface thermal proteome profiling tracks perturbations and drug targets on the plasma membrane. Nat. Methods 18, 84–91 (2021).

30. Heterologous Gene Expression in E.Coli: Methods and Protocols. vol. 1586 (Springer New York, New York, NY, 2017).

31. Barnaba, C. et al. Lipid-exchange in nanodiscs discloses membrane boundaries of cytochrome-P450 reductase. Chem. Commun. 54, 6336–6339 (2018).

32. Van Meer, G., Voelker, D. R. & Feigenson, G. W. Membrane lipids: where they are and how they behave. Nat. Rev. Mol. Cell Biol. 9, 112–124 (2008).

33. Brignac-Huber, L. M., Park, J. W., Reed, J. R. & Backes, W. L. Cytochrome P450 Organization and Function Are Modulated by Endoplasmic Reticulum Phospholipid Heterogeneity. Drug Metab. Dispos. 44, 1859–1866 (2016).

34. Babu, M. et al. Global landscape of cell envelope protein complexes in Escherichia coli. Nat. Biotechnol. 36, 103–112 (2018).

35. Jandu, R. S. et al. Capture of endogenous lipids in peptidiscs and effect on protein stability and activity. iScience 27, 109382 (2024).

36. Chen, Y. & Duong Van Hoa, F. Peptidisc-Assisted Hydrophobic Clustering Toward the Production of Multimeric and Multispecific Nanobody Proteins. Biochemistry 64, 655–665 (2025).

37. Johnston, H. E. et al. Solvent Precipitation SP3 (SP4) Enhances Recovery for Proteomics Sample Preparation without Magnetic Beads. Anal. Chem. 94, 10320–10328 (2022).

38. Cox, J. & Mann, M. MaxQuant enables high peptide identification rates, individualized p.p.b.-range mass accuracies and proteome-wide protein quantification. Nat. Biotechnol. 26, 1367–1372 (2008).

39. Cox, J. et al. Accurate Proteome-wide Label-free Quantification by Delayed Normalization and Maximal Peptide Ratio Extraction, Termed MaxLFQ. Mol. Cell. Proteomics 13, 2513–2526 (2014).

40. Demichev, V., Messner, C. B., Vernardis, S. I., Lilley, K. S. & Ralser, M. DIA-NN: neural networks and interference correction enable deep proteome coverage in high throughput. Nat. Methods 17, 41–44 (2020).

41. Tyanova, S. et al. The Perseus computational platform for comprehensive analysis of (prote)omics data. Nat. Methods 13, 731–740 (2016).

42. Wickham, H. et al. Welcome to the Tidyverse. J. Open Source Softw. 4, 1686 (2019).

43. Wu, T., et al. clusterProfiler 4.0: A universal enrichment tool for interpreting omics data. The Innovation 2, 100141 (2021).

44. Huber, W. et al. Orchestrating high-throughput genomic analysis with Bioconductor. Nat. Methods 12, 115–121 (2015).

45. Käll, L., Krogh, A. & Sonnhammer, E. L. L. A Combined Transmembrane Topology and Signal Peptide Prediction Method. J. Mol. Biol. 338, 1027–1036 (2004).

46. The UniProt Consortium et al. UniProt: the Universal Protein Knowledgebase in 2025. Nucleic Acids Res. 53, D609–D617 (2025).

47. Schneider, C. A., Rasband, W. S. & Eliceiri, K. W. NIH Image to ImageJ: 25 years of image analysis. Nat. Methods 9, 671–675 (2012).

48. Perez-Riverol, Y. et al. The PRIDE database at 20 years: 2025 update. Nucleic Acids Res. 53, D543–D553 (2025).

